# Forward genetics in *Wolbachia*: Regulation of *Wolbachia* proliferation by the amplification and deletion of an addictive genomic island

**DOI:** 10.1101/2020.09.08.288217

**Authors:** Elves H Duarte, Ana Carvalho, Sergio López-Madrigal, João Costa, Luís Teixeira

**Affiliations:** Instituto Gulbenkian de Ciência, Oeiras, Portugal.; Faculdade de Ciências e Tecnologia, Universidade de Cabo Verde, Palmarejo, Cabo Verde.; Department of Genetics, University of Cambridge, Cambridge, United Kingdom.; Department of Evolutionary Ecology, Institute of Organismic and Molecular Evolution, Johannes Gutenberg University, Mainz, Germany; Faculdade de Medicina, Universidade de Lisboa, Lisboa, Portugal.

**Author notes:** corresponding authors: Elves H Duarte Luís Teixeira.

## Abstract

*Wolbachia* is one of the most prevalent bacterial endosymbionts, infecting approximately 40% of terrestrial arthropod species. *Wolbachia* is often a reproductive parasite but can also provide fitness benefits to its host, as, for example, protection against viral pathogens. This protective effect is currently being applied to fight arboviruses transmission by releasing *Wolbachia*-transinfected mosquitoes. Titre regulation is a crucial aspect of *Wolbachia* biology. Higher titres can lead to stronger phenotypes and fidelity of transmission but can have a higher cost to the host. Since *Wolbachi*a is maternally transmitted, its fitness depends on host fitness, and, therefore, its cost to the host may be under selection. Understanding how *Wolbachia* titres are regulated and other aspects of *Wolbachi*a biology has been hampered by the lack of genetic tools. Here we developed a forward genetic screen to identify new *Wolbachia* over-proliferative mutant variants. We characterized in detail two new mutants, *w*MelPop2 and *w*MelOctoless, and show that the amplification or loss of the Octomom genomic region lead to over-proliferation. These results confirm previous data and expand on the complex role of this genomic region in the control of *Wolbachia* proliferation. Both new mutants shorten the host lifespan and increase antiviral protection. Moreover, we show that *Wolbachia* proliferation rate in *Drosophila melanogaster* depends on the interaction between Octomom copy number, the host developmental stage, and temperature. Our analysis also suggests that the life shortening and antiviral protection phenotypes of *Wolbachia* are dependent on different, but related, properties of the endosymbiont; the rate of proliferation and the titres near the time of infection, respectively. Altogether, we demonstrate the feasibility of a novel and unbiased experimental approach to study *Wolbachia* biology, which can be further adapted to characterize other genetically intractable bacterial endosymbionts.

## Introduction

Intracellular maternally-transmitted bacterial symbionts (bacterial endosymbionts) are widespread in insects [1]. These bacterial endosymbionts can be mutualistic by, for instance, complementing the diets of their hosts, and may expand the range of ecological niches of their insect hosts [1]. They can also be parasitic, often manipulating the reproduction of their hosts and promoting their spread in the host population [1]. Understanding the interaction of endosymbionts with their hosts is crucial to understand much of insect biology. A key aspect of this interaction is the regulation of endosymbiont titres, which influence the strength of the induced phenotypes and the cost to the hosts [2,3].

*Wolbachia* is one of the most prevalent bacterial endosymbionts in arthropods, being found in approximately 40% of terrestrial arthropod species [4]. *Wolbachia* is broadly known as a host reproduction manipulator [5]. However it can also be mutualistic, by, for example, vitamin provision [6] or protection against viral pathogens [7,8].

The discovery of *Wolbachia*-induced protection against viruses in *Drosophila melanogaster*, prompted its use to control arboviruses transmission by insect vectors [9]. *Aedes aegypti* mosquitoes trans-infected with *Wolbachia* have increased resistance to viruses, including dengue, chikungunya, Zika, and yellow fever viruses, and, therefore, reduced vector competence [10–13]. Release of *Wolbachia*-infected mosquitoes in dengue-endemic areas is likely to reduce dengue burden [14,15]. Despite the preliminary successful results of this strategy, we still lack knowledge on several fundamental aspects of *Wolbachia* biology and interaction with viral pathogens, which hinders predicting the long-term outcome of *Wolbachia*-based interventions to control insect-vector transmitted viruses.

*Wolbachia* titres are a critical factor regulating its biology and interaction with the host [3]. Titres correlate positively with transmission fidelity and the strength of *Wolbachia*-induced phenotypes, including the *Wolbachia* pathogen blocking phenotype [3,16–20]. In contrast, higher titres are associated with a reduction in host lifespan [16,17,21,22]. This may also have a cost to *Wolbachia*, since as a vertically transmitted bacterium, its fitness depends on the host fitness. Thus, *Wolbachia* titres regulation by the symbiont or the host may be under selection. Although several host and environmental factors (e.g. temperature) have been shown to affect *Wolbachia* titres, less is known about *Wolbachia* genes that regulate its titres [3].

So far, a single *Wolbachia* genetic factor, the Octomom region, has been shown to influence proliferation [16,17]. This genomic region, predicted to encode eight genes, is amplified in the highly proliferative and pathogenic *w*MelPop. Moreover, the degree of amplification of the Octomom region determines the proliferation rate of *w*MelPop and the strength of its life shortening phenotype [17].

The genetic intractability of *Wolbachia*, which remains unculturable so far, hampers the identification of more genetic modifications altering *Wolbachia* proliferation. Hence, unbiased approaches such as genetic screens could contribute to our understanding of the genetic bases of *Wolbachia*-host interactions. Here, we developed a screening strategy in *Wolbachia* to isolate novel over-proliferating variants. The strategy was based on random mutagenesis, which has been applied before to other unculturable bacteria [23]. We fed the mutagen ethyl methanesulfonate (EMS) to *D. melanogaster* females carrying *Wolbachia* and screened for over-proliferative *Wolbachia* in their progeny. This approach allowed us to isolate new mutant over-proliferating *Wolbachia* variants. We identified the causative genetic changes in *Wolbachia* causing over-proliferation and made a detailed phenotypical characterization in terms of proliferation, cost to the host, and antiviral protection. We identified a new mutation leading to *Wolbachia* over-proliferation and revealed a complex role for the Octomom region in regulating *Wolbachia* proliferation. Moreover, we demonstrated the feasibility of a novel and unbiased experimental approach to study *Wolbachia* biology.

## Results

### Isolation of over-proliferative *Wolbachia* in an unbiased forward genetic screen

We implemented a classical forward genetic screen in order to isolate new over-proliferative *Wolbachia* variants. We attempted to mutagenize *Wolbachia* by feeding the mutagen EMS to *Wolbachia*-carrying *D. melanogaster* females. EMS is extensively used in *D. melanogaster* [24] and has been previously used to mutagenize intracellular bacteria in cell culture [23]. We then tested *Wolbachia* titres, by real-time quantitative PCR (qPCR), in the progeny of treated females, since this bacterium is maternally transmitted. We used flies with the variant *w*MelCS_b as our starting variant because of its potential to easily become over-proliferative, given its genetic proximity to the over-proliferative and pathogenic *w*MelPop variant [16,17,22,25].

Putative mutagenized *Wolbachia* cells within the host would be in a mixed population, which would make it harder to assess their specific phenotype. However, we hypothesized that over-proliferating *Wolbachia* cells could overtake the population and that the resulting higher titres could be detectable. Moreover, we decided to pre-treat some of the EMS exposed females with tetracycline to reduce the *Wolbachia* population in these females and their progeny. This *Wolbachia* titre reduction should decrease competition for any new mutated *Wolbachia*, increase drift during vertical transmission, and, therefore, potentially facilitate fixation of new variants. To set up the conditions for tetracycline treatment, we tested different doses of this antibiotic on females, without EMS. The progeny of treated females had from 0 to 90% of the *Wolbachia* titres in controls (S1 Fig, *p* < 0.001 for all doses compared with control, at generation 1). We then followed the subsequent progeny of these flies to test how many fly generations it takes to recover normal *Wolbachia* titres. Except for higher tetracycline doses which lead to infection loss, *Wolbachia* titres recovered to normal within four fly generations (S1 Fig; linear mixed model [lmm], *p* > 0.48 for all doses compared with control at generation 4).

We also tested for the effect of different EMS doses on the fecundity of *D. melanogaster* females and *Wolbachia* titres. We observed that increasing doses of EMS reduce female fecundity (S2A-B Fig, linear model [lm], *p* < 0.001 for both egg number and adult progeny per female). Moreover, we found that EMS feeding strongly reduces *Wolbachia* titres in the next generation, in a dose-dependent manner (S2C-D Fig, non-linear model [nls] fit, *p* < 0.001). Titres were reduced by up to 90% when 8,000 mM EMS was supplied, leading to the loss of *Wolbachia* in the next generation in some lines (S2C-D Fig). Given these results and the recovery time after tetracycline treatment detailed above, we quantified *Wolbachia* titres at the first generation (F1), the immediate progeny of EMS-treated females, and at the fourth generation after treatment (F4), when we would expect *Wolbachia* titres to recover after the severe reduction due to EMS treatment.

We screened approximately one thousand F1 progeny of EMS-treated females, in a range of experimental conditions, and at least one F4 female descendent per treated female. We varied EMS dose from 10 mM to 8,000 mM, and tetracycline dose from 0 μg/ml to 12.5 μg/ml, in different combinations (S1 Table). The relative *Wolbachia* titre was determined when females were ten days old, after they laid eggs, so that any putatively interesting progeny could be followed up.

In three independent batches of EMS-treated flies, we detected females with 3 to 14-fold more *Wolbachia* than controls, suggesting the presence of over-proliferative variants (Fig 1 and S3 Fig). In two batches, over-proliferating *Wolbachia* were identified in the F1 and in the other batch in the F4. We assessed *Wolbachia* titres in the next generation and found that the over-proliferative phenotypes were inherited. Subsequent selection allowed us to establish *D. melanogaster* lines carrying new potentially over-proliferative *Wolbachia* variants.

**Fig 1.**
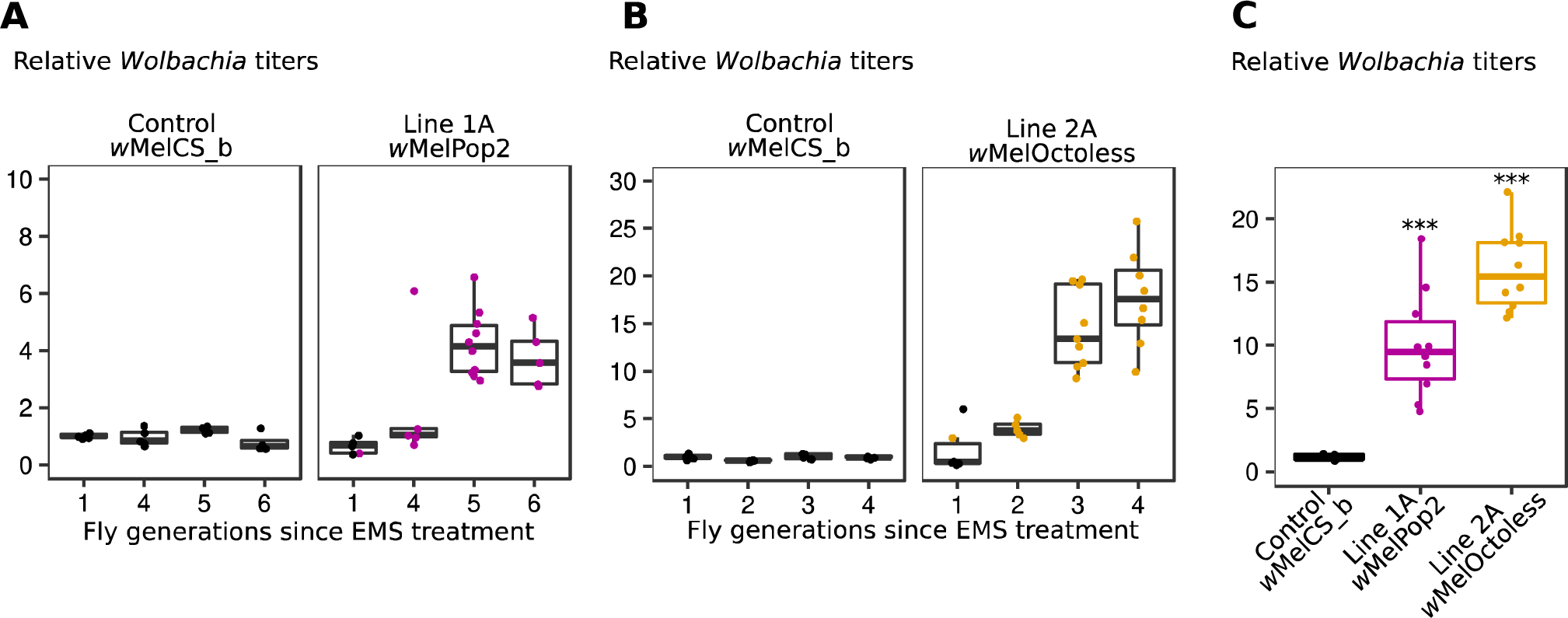
Isolation of over-proliferative *Wolbachia* variants by a forward genetic screen. (A and B) Relative *Wolbachia* titres in a control (*w*MelCS_b) and EMS-treated flies (Lines 1A and 2A). 5–10 virgin females were randomly collected each generation for egg-laying and *Wolbachia* titre measurement using qPCR. Bacterial titres are normalized to that of control flies. The female used in the first generation to start the next generation is coloured. At the other generations the progeny of the female with the higher *Wolbachia* titre was used to set up the next generation. The selection of the other putative over-proliferating *Wolbachia* line in panel B in shown in S3A Fig. (C) Relative titres of over-proliferating *Wolbachia* variants in a host isogenic genetic background. Both lines kept the over-proliferative phenotype (p < 0.001). Each dot represents the *Wolbachia* titre of a single female.

We designed the screen to find new mutants of *Wolbachia* that lead to the endosymbiont over-proliferation. However, EMS will most likely also induce mutations in the host, in the nuclear or mitochondrial genomes, that can be transmitted. To minimize the influence of host nuclear mutations on our screen, we backcrossed the EMS-treated females and their progeny, at every generation, with males from the control isogenic line. To verify that new mutations in the host were not the cause of *Wolbachia* over-proliferation, we replaced the first, second and third chromosomes of *D. melanogaster* females carrying the over-proliferating *Wolbachia* variants in lines 1A, 2A, and 3A, with the chromosomes of the control line, through the use of balancer chromosomes (S4 Fig). We then repeated *Wolbachia* titres quantification and found that the over-proliferative phenotypes were maintained (Fig 1C, S5 Fig; lmm, p-value< 0.001 for all compared with *w*MelCS_b).

Since mitochondria are maternally transmitted and could have been also mutated by EMS, the experiments described above cannot exclude the possibility that *Wolbachia* over-proliferation is mitochondria-determined. Thus, the mitogenome of the lines 1A and 2A, showing higher *Wolbachia* titres, were sequenced with Illumina short-reads and aligned to the mitochondrial reference genome release 6 (GenBank: KJ947872.2:1–14,000, S2 Table). We did not find SNPs or indels unique to the mitochondria of these flies, which shows that flies with over-proliferative *Wolbachia* did not inherit mutated mitochondria (S3 Table). Therefore, we concluded that the observed *Wolbachia* over-proliferative phenotypes did not result from mutations in neither the nuclear or mitochondrial host genome.

### Identification of genetic basis of the new over-proliferative variants

To identify the mutations associated with over-proliferation, we sequenced and assembled the genomes of these over-proliferative *Wolbachia*. We performed a hybrid assembly with short (Illumina) and long-reads (Nanopore) and obtained single and circular genomes for each *Wolbachia* chromosome (S2 and S4 Tables).

To test our assembly pipeline we sequenced and assembled a previously characterized Cluster III *w*Mel *Wolbachia* variant, named *w*Mel [16], which derives from the line used for the original *w*Mel reference genome (GenBank: AE017196.1) [26]. The new *w*Mel genome (GenBank: CP046925.1) was also circular and comparable in size, structure and number of ORFs with previously published *w*Mel genomes [26,27], including the *w*Mel reference genome (S4 Table). We found, however, two SNPs and seven indels relative to the *w*Mel reference genome, which we confirmed using Sanger sequencing (S5 Table). These results validate our sequencing pipeline.

We assembled the genome of the control variant *w*MelCS_b using this pipeline (GenBank: CP046924.1), in order to be able to identify new mutations in the new variants. We also compared this new assembly of *w*MelCS_b with the *w*Mel reference genome (GenBank: AE017196.1) and identified 37 indels and 146 SNPs between these variants (S6 Table).

The only difference between the genome of the over-proliferative *Wolbachia* variant in Line 1A (GenBank: CP046922.1) and *w*MelCS_b was an amplification of the Octomom region (Fig 2A and S1 Text). There were three more copies of this region, giving a genome size difference of 62,814bp. The Octomom region amplification, and lack of other differences, was also confirmed by mapping of the Illumina sequencing reads from Line 1A on the genome of *w*MelCS_b (GenBank: CP046924.1) and by qPCR (Fig 2B and S6 Fig). These results show that Octomom amplification is the cause of over-proliferation, consistently with previous findings with the variant *w*MelPop [16,17,28]. As shown before for *w*MelPop [17], we observed variation in the Octomom copy number in *w*MelPop2-carrying flies. For further analyses of the phenotype of this variant we established, through selection (as in [17]), *D. melanogaster* lines carrying *Wolbachia* with low (2-3) or high Octomom (8-9) copy number (S7 Fig). We named this variant *w*MelPop2, given the nature of the genomic change inducing its over-proliferation.

**Fig 2.**
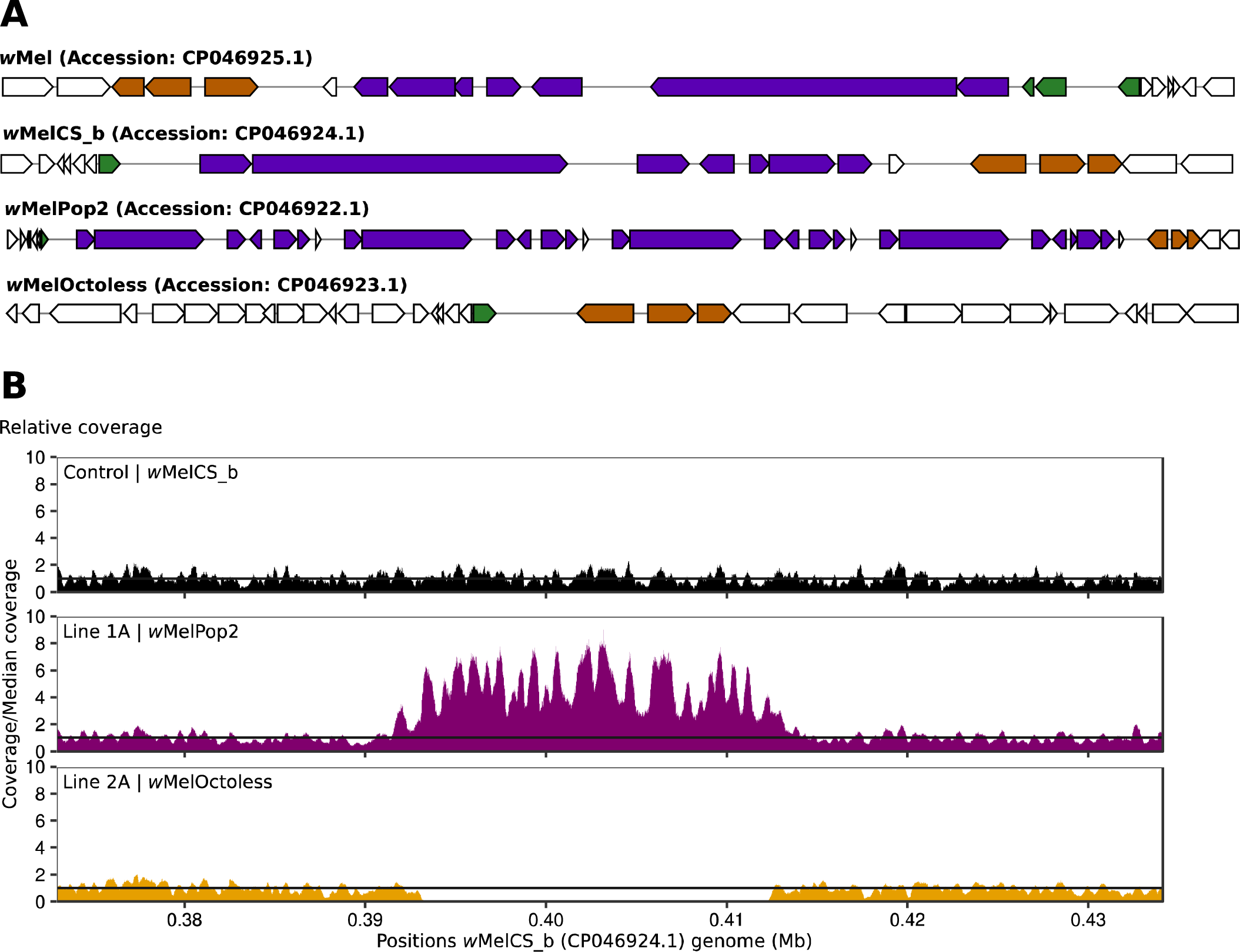
Both amplification and deletion of Octomom lead to *Wolbachia* over-proliferation. (A) Representation of Octomom region and its flanking region in over-proliferative *Wolbachia*. *De novo* assembled genomes of *w*Mel, *w*MelCS_b, Line1A (*w*MelPop2) and Line 2A (*w*MelOctoless) were annotated using the NCBI Prokaryotic Genome Annotation Pipeline v4.10. These representations were generated using MultiGeneBlast v1.1.13 (http://multigeneblast.sourceforge.net/) and identify the homologous genes immediately upstream of Octomom (orange), in the Octomom region (purple), and immediately downstream of Octomom (green), in the *w*Mel reference genome (GenBank: AE017196.1). Regions are not to scale. Note that the genome annotation differs between this new *w*Mel genome assembly and the reference genome, although there is no difference between the sequences in this region. (B) Relative coverage in the genomic region containing the Octomom region. Illumina paired-end reads of the different *Wolbachia* variants were mapped to *w*MelCS_b genome (GenBank: CP046924.1).

We sequenced and assembled the *w*MelPop genome following the same pipeline (GenBank: CP046921.1) and compared it to *w*MelPop2. We only detected the two SNPs previously identified between *w*MelCS_b and *w*MelPop (position 920,191: T in *w*MelPop and C in *w*MelPop2; and position 1,005,339: A in *w*MelPop and G in *w*MelPop2 [16]). We also compared the mitogenome of flies carrying *w*MelPop and *w*MelPop2 and found one single substitution (position 10,793: G in WMelPop and A in wMelPop2) (S3 Table), which we confirmed using Sanger sequencing. Genome assembly and individual Nanopore long-reads from *w*MelPop and *w*MelPop2 (S7 Table, S8 Fig) show that Octomom amplification in these variants occurs in tandem, as previous data indicated [17].

Interestingly, the genome of the over-proliferative *Wolbachia* variant in line 2A (GenBank: CP046923.1) only differs from *w*MelCS_b by a deletion of a 20,938bp genomic fragment that includes the full-length Octomom region and one of its flaking direct repeats (Fig 2A and S1 Text). Mapping the Illumina sequencing reads of this variant on the genome of *w*MelCS_b (GenBank: CP046924.1) confirmed this deletion as the only difference between the two (Fig 2B). The absence of all Octomom genes in this line was also confirmed by qPCR (S6 Fig). These results identify loss of the Octomom region as the cause of this variant over-proliferative phenotype. Thus, we named this variant *w*MelOctoless.

The variant in line 2B, isolated together with *w*MelOctoless, also lost the Octomom region. This was the only observed difference when mapping the Illumina reads on *w*MelCS_b (S9 Fig), and no differences were observed when the Illumina reads were mapped to the *w*MelOctoless genome (GenBank: CP046923.1). Since *w*MelOctoless and the variant in line 2B were identified in the same batch of mutagenesis, they may be not independent. However, and importantly, we obtained the same results with another independent over-proliferative line, isolated in a different batch of treatment, line 3A (S3 and S9 Fig). Mapping the Illumina sequence reads from this line to *w*MelCS_b also identifies the loss of Octomom as the only mutation in this variant. Accordingly, there are no differences to *w*MelOctoless. Therefore, we named this line *w*MelOctoless2. These results further confirm that loss of the Octomom region leads to an over-proliferative phenotype in *Wolbachia*.

In summary, we were able to identify the genomic changes associated with the new over-proliferative variants and all map to loss or amplification of the Octomom region.

### Deletion and amplification of the Octomom region differently impact titres and growth of *Wolbachia*

In order to characterize better the phenotypes of the new *Wolbachia* variants *w*MelOctoless and *w*MelPop2, we analysed their proliferation, together with *w*MelCS_b and *w*MelPop, in adult males kept at 18°C, 25°C, and 29°C (Fig 3 and S10 Fig). The flies were reared at 25°C and placed at the different temperatures when 0-1 days old adults. At this initial point, at adult eclosion, there are differences in titres between lines carrying different *Wolbachia* variants (S11 Fig, *p* < 0.028 for all comparisons). Flies carrying *w*MelCS_b have the lowest relative titre of *Wolbachia.* Flies carrying variants with low amplification of the Octomom region have approximately twice the titres of *Wolbachia*, while flies carrying variants with high copy number of this region have three times more *Wolbachia* than *w*MelCS_b. Finally, flies carrying *w*MelOctoless have the highest titres, approximately four-fold higher than flies carrying *w*MelCS_b. Therefore, the deletion or amplification of the Octomom region impact *Wolbachia* titres at adult eclosion.

**Fig 3.**
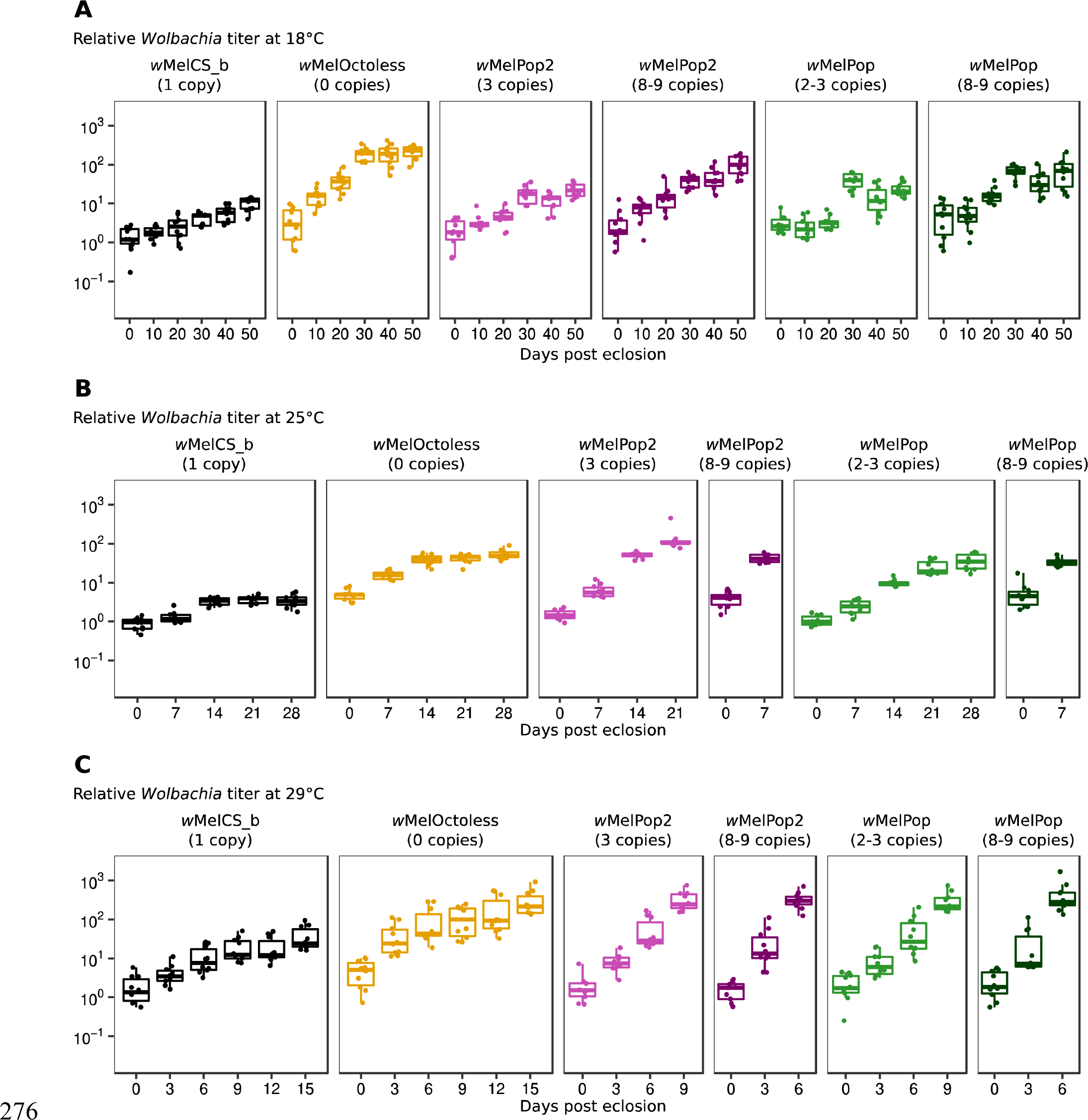
The amplification or deletion of Octomom increase *Wolbachia* proliferation rate in adults. Time-course of relative *Wolbachia* titres in adults at 18°C (A), 25°C (B) and 29°C (C) with different *Wolbachia* variants. *D. melanogaster* males used in these experiments developed at 25°C, were collected on the day of adult eclosion and aged at the given temperatures (18°C, 25°C or 29°C). Ten males were collected at each time-point for *Wolbachia* titre measurement using qPCR. *Wolbachia* titres were normalized to that of 0-1 days-old *w*MelCS_b-infected males. A replicate of the experiment is shown in S10 Fig. Exponential models were used to estimate *Wolbachia* doubling time, using both full replicates, and a summary of the results is given in Table 1. Each dot represents the relative *Wolbachia* titre of a single male.

To analyse proliferation during adult life, we fitted an exponential model to the titres over adult age and estimated doubling time of the *Wolbachia* variants, at different temperatures (Table 1). *Wolbachia* doubling time varies widely with *Wolbachia* variant and temperature, from approximately one day to seventeen days. A model with all the data shows a complex interaction between proliferation, *Wolbachia* variant and temperature (lmm, *p* < 0.001). We analysed this dataset by comparing specific set of variants to test differences between *w*MelOctoless and *w*MelCS_b, differences between *w*MelPop2 and *w*MelCS_b, and differences between levels of Octomom amplification in *w*MelPop and *w*MelPop2.

**Table 1.** Doubling time of *Wolbachia* variants in larvae and adults at different temperatures.

|  | Doubling time - days (95% confidence interval) |  |  |  |
| --- | --- | --- | --- | --- |
|  | Larvae | Adults |  |  |
|  | 25°C | 18°C | 25°C | 29°C |
| wMelCS_b | 0.68<br>(0.55-0.91) | 16.79<br>(13.29-22.79) | 13.90<br>(9.84-23.62) | 4.07<br>(3.35-5.17) |
| wMelOctoless | 0.50<br>(0.42-0.61) | 9.41<br>(8.20-11.04) | 10.46<br>(7.99-15.16) | 3.37<br>(2.86-4.10) |
| wMelPop2 (3 copies) | n.d. | 17.68<br>(13.84-24.45) | 3.50<br>(3.02-4.15) | 1.28<br>(1.13-1.48) |
| wMelPop2 (8-9 copies) | 0.59<br>(0.48-0.75) | 11.86<br>(10.00-14.56) | 2.26<br>(1.74-3.22) | 1.07<br>(0.92-1.28) |
| wMelPop (2-3 copies) | n.d. | 15.44<br>(12.43-20.36) | 5.30<br>(4.50-6.44) | 1.38<br>(1.22-1.60) |
| wMelPop (8-9 copies) | n.d. | 13.82<br>(11.33-17.70) | 2.79<br>(2.04-4.43) | 0.88<br>(0.76-1.02) |

n.d. – not determined

A direct comparison between *w*MelOctoless with *w*MelCS_b shows that this new variant replicates faster than *w*MelCS_b (lmm, *p* < 0.001), although it is a relatively small difference at all temperatures (in the full model with all variants, however, the proliferation of *w*MelOctoless and *w*MelCS_b is only statistically different at 18°C, Table 1). Both strains interact equally with temperature. Their growth rate does not significantly change between 18°C and 25°C (*p* = 0.94), but increases at 29°C (*p* < 0.001).

A comparison of *w*MelPop2 having high and low Octomom copy number with *w*MelCS_b and *w*MelOctoless shows that these variants with Octomom amplification have the highest growth rates at 25°C and 29°C (*p* < 0.001 for all comparisons of *w*MelPop2 (3 or 8-9 copies) compared to *w*MelCS_b and *w*MelOctoless). At 18°C *w*MelPop2 with 3 copies of Octomom has a growth rate similar to *w*MelCS_b (*p* = 0.79) and lower than *w*MelOctoless (*p* < 0.001). While at this temperature the growth rate of *w*MelPop2 with 8-9 copies of Octomom is not significantly different from either *w*MelCS_b or *w*MelOctoless (*p* > 0.088 in both comparisons), and the estimated value is in-between the two (Table 1). The analysis also shows a strong interaction between *w*MelPop2 growth and temperature. Both low and high Octomom copy number *w*MelPop2 growth rates increase from 18°C to 25°C, and from 25°C to 29°C (*p* < 0.001 for these comparisons).

To test the effect of the degree of Octomom amplification on growth rate and differences between *w*MelPop and *w*MelPop2, we compared these variants with low or high copy number of the Octomom region. The variants with the high copy number have a higher growth rate than the variants with low copy number at all temperatures (*p* < 0.025 at all temperatures). These results confirm that the degree of amplification of the Octomom region controls the intensity of the over-proliferation of these variants, as shown before [17]. Both low and high Octomom copy number *w*MelPop and *w*MelPop2 increase growth rate with temperature (*p* < 0.001 for low and high copy number variants compared between 18°C and 25°C, and between 25°C and 29°C), confirming the analysis above.

The statistical model comparing *w*MelPop and *w*MelPop2, which differ in two SNPs (see above), indicated a significant difference in growth between them at 25°C (*p* < 0.001). This could indicate that these two SNPs also influence growth of *Wolbachi*a. However, this could also be due to the fact that the copy number of the Octomom region was not equally controlled in *w*MelPop and *w*MelPop2 lines during these experiments. *w*MelPop low copy number line had 2-3 copies of Octomom, while the *w*MelPop2 line had 3 copies. To test if *w*MelPop and *w*MelPop2 indeed vary in proliferation rate, we repeated this experiment with a more tightly controlled Octomom copy number in these two lines, at 25°C (S12 Fig A-B). Both *w*MelPop and *w*MelPop2 carrying 3 copies of Octomom grow faster than *w*MelCS_b (lmm, *p* < 0.001 for both) and there is no difference in growth between them (*p* = 0.39). This indicates that the genetic differences between these lines do not affect their growth and that they are equally influenced by Octomom copy number.

Overall, the data and analysis show a complex interaction between *Wolbachia* variants, temperature and growth rate. There is a strong interaction between temperature and the increased proliferation of variants with amplification of the Octomom region, *w*MelPop and *w*MelPop2, when compared with *w*MelCS_b. The effect of the amplification is not significant at 18°C and becomes increasingly stronger at higher temperatures. On the other hand, loss of Octomom leads to a smaller effect in growth, but similar at all temperatures, when compared with *w*MelCS_b. Therefore, although both genomic mutations lead to an increase in *Wolbachia* titres they have different impacts in the growth rates and interaction with temperature.

### Rapid proliferation of *Wolbachia* during the host development

We also analysed the growth of *w*MelCS_b, *w*MelOctoless and *w*MelPop2 (8-9 Octomom copies) during host development. *D. melanogaster* develops from egg to adult in only 10 days, at 25°C. We predicted that *Wolbachi*a would grow much faster during this period than during adult life, considering changes in *Wolbachia* loads from eggs to adults [29]. We, therefore, estimated absolute numbers of *Wolbachia* genome copies in individuals at the different stages of development using qPCR for the single copy gene *wsp* and a calibration curve using a plasmid with *wsp* cloned. Assuming one chromosome per *Wolbachia* cell [30], these numbers correspond to *Wolbachia* cells. Embryos with 0-2h have between 2,300 and 3,100 *Wolbachia* genome copies, with no significant difference between *Wolbachia* variants (lm, *p* = 0.87 for the effect of *Wolbachia* variant, Fig 4, Table 2). At the end of development, newly eclosed adults carry from approximately 400,000 to 3,200,000 *Wolbachia* genome copies. At this stage, however, and as observed above (S11 Fig), there are significant differences between the three variants (lm, *p* < 0.008 for all comparisons, Table 2, Fig 4). Also, males carry less *Wolbachia* than females (*p* = 0.033).

**Fig 4.**
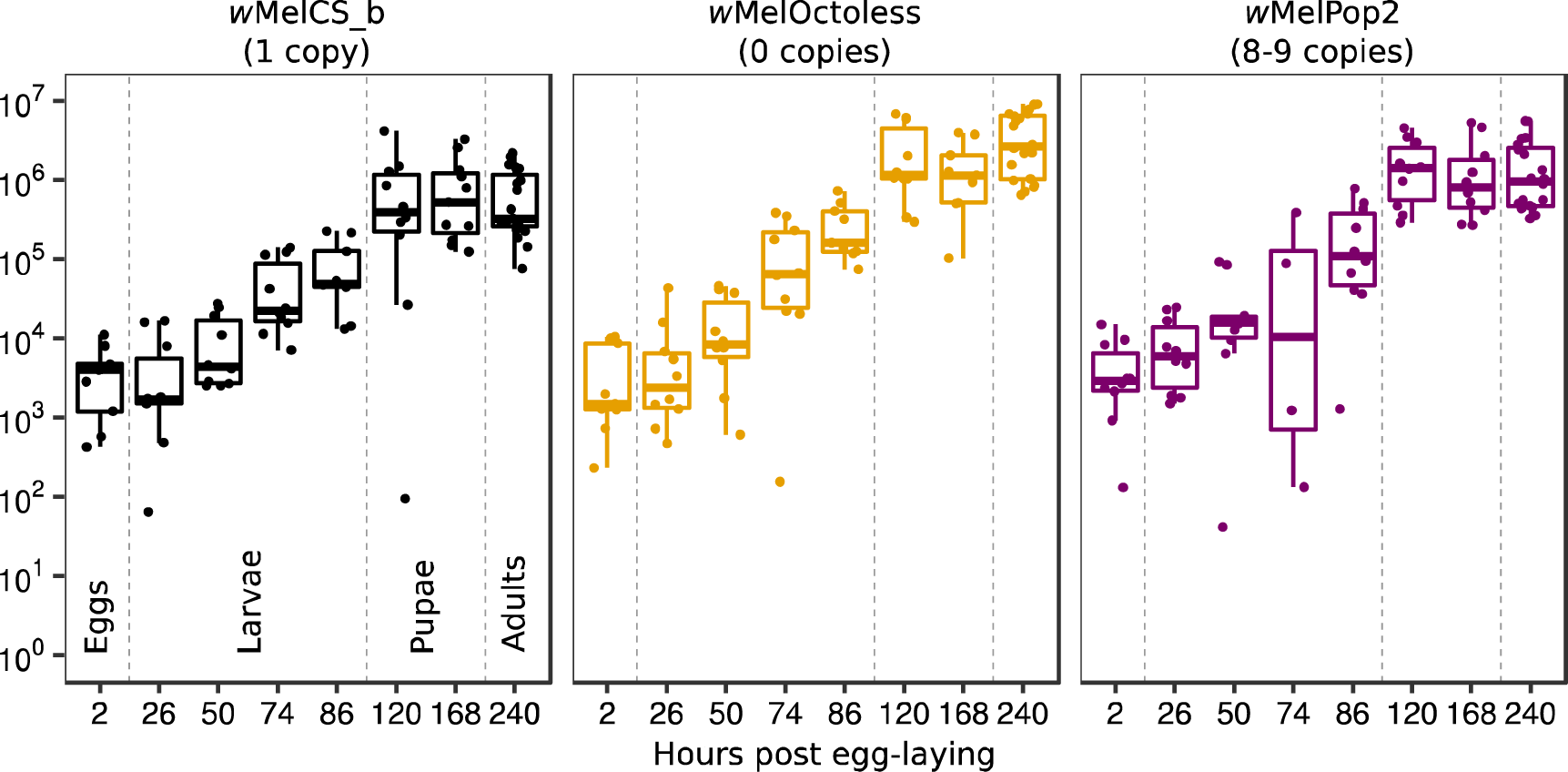
*Wolbachia* proliferates rapidly during larval development. *Wolbachia* genome copies throughout *D. melanogaster* development. Samples are embryos (2h), 1^st^ instar larvae (26h), 2^nd^ instar larvae (50h), 3^rd^ instar larvae (74h and 86h), white prepupae (120h), P8 staged pupae (168h), and newly eclosed adults (240h). Vertical dashed lines separate developmental stages (i.e. eggs, larvae, pupae, and adults). Dots represent either a pool of 10 individuals (embryos and larvae) or a single individual (pupae and adults). The x-axis is not in a continuous scale. Proliferation of the different *Wolbachia* variants in the first 120 hours was analysed using an exponential model. A summary is given in Table 1.

**Table 2.**
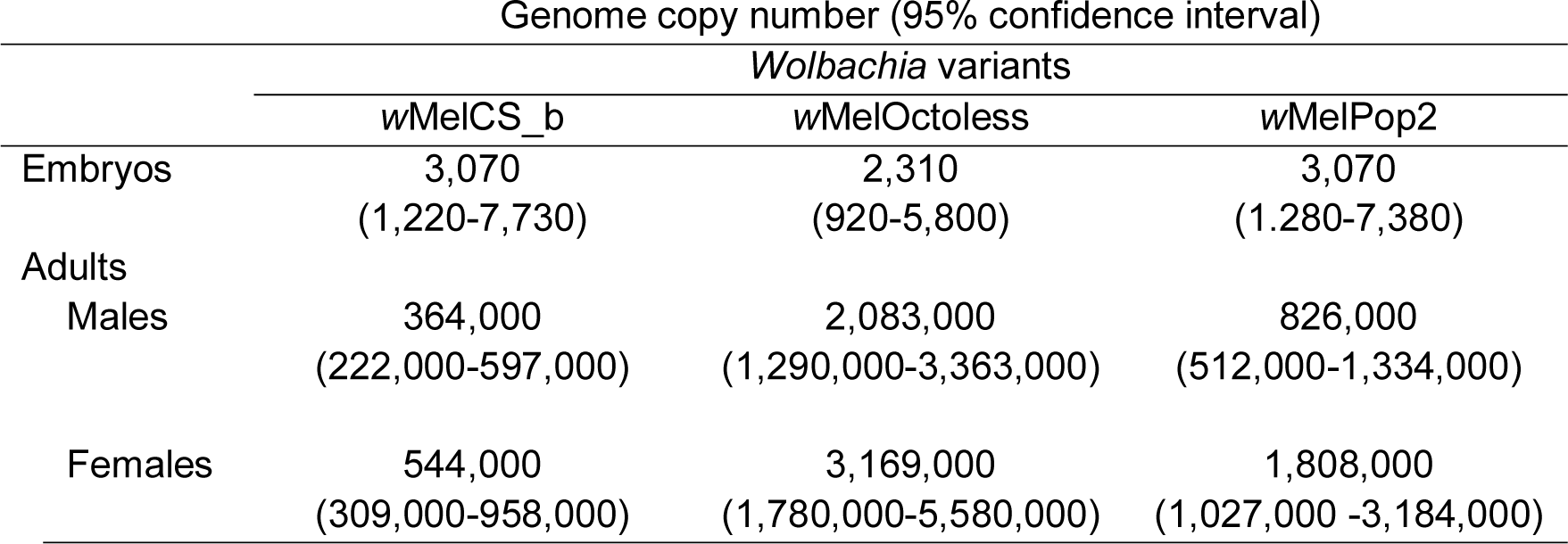
*Wolbachia* genome copies in embryos and newly eclosed adult flies.

*Wolbachia* growth seems to be restricted to the period between egg and white prepupae (120h), since there is no significant growth from this stage to adults (lm, *p* = 0.46). From eggs to white prepupae there is rapid exponential growth of all variants (Fig 4, Table 1). *w*MelCS_b has an estimated doubling time of approximately 16h, *w*MelPop2 of 14h, and *w*MelOctoless of 12h. These different doubling times probably explain how *Wolbachia* reach different amounts per individual host in adults, starting from the same estimated amount in embryos. However, in this analysis the difference between growth rates is not statistically significant (*p* = 0.12 for interaction between *Wolbachia* variants and growth).

The growth rates of these variants are, therefore, very similar during this stage, and much faster than in adults. At the same temperature, we estimated doubling times in adults of *w*MelCS_b, *w*MelOctoless, and *w*MelPop2 (high-copy) to be, approximately, 13.9, 10.5, and 2.3 days, respectively (Table 1). Therefore, *Wolbachia* growth at different stages of *D. melanogaster* can vary dramatically, and the different variants respond differently to different stages of the host life cycle.

We also asked if *Wolbachia* Octomom copy number changed in *w*MelCS_b and *w*MelPop2, throughout development, as *Wolbachia* is proliferating fast, and found no evidence of so (lm, *p* = 0.49, S13A Fig). However, during adult life there was a small increase of Octomom copy number with age in *w*MelPop and *w*MelPop2 (an increase of 0.032 per day, lmer, *p* = 0.009, S13B Fig), as shown before [31].

### *Wolbachia* variants with a deletion or amplification of the Octomom region induce different life-shortening phenotypes

The over-proliferation of *w*MelPop has been associated with a shortening of the host lifespan [16,22]. We, therefore, tested if these new over-proliferative variants also shorten the lifespan of their host, at different temperatures, in males (Fig 5A-D, S14A-C Fig). We also performed this assay in females at 25°C, with similar results to males at 25°C (S14D-E Fig). There was a significant interaction between *Wolbachia* variant and temperature (Cox proportional hazard model with mixed effects (CHR), *p* < 0.001). All lines, including the *Wolbachia*-free line have a shorter lifespan at 25°C than at 18°C, and even shorter at 29°C (*p* < 0.001 for all these comparisons). *w*MelCS_b did not affect the host lifespan at any temperature (*p* > 0.16 for all comparisons with the *Wolbachia*-free line).

**Fig 5.**
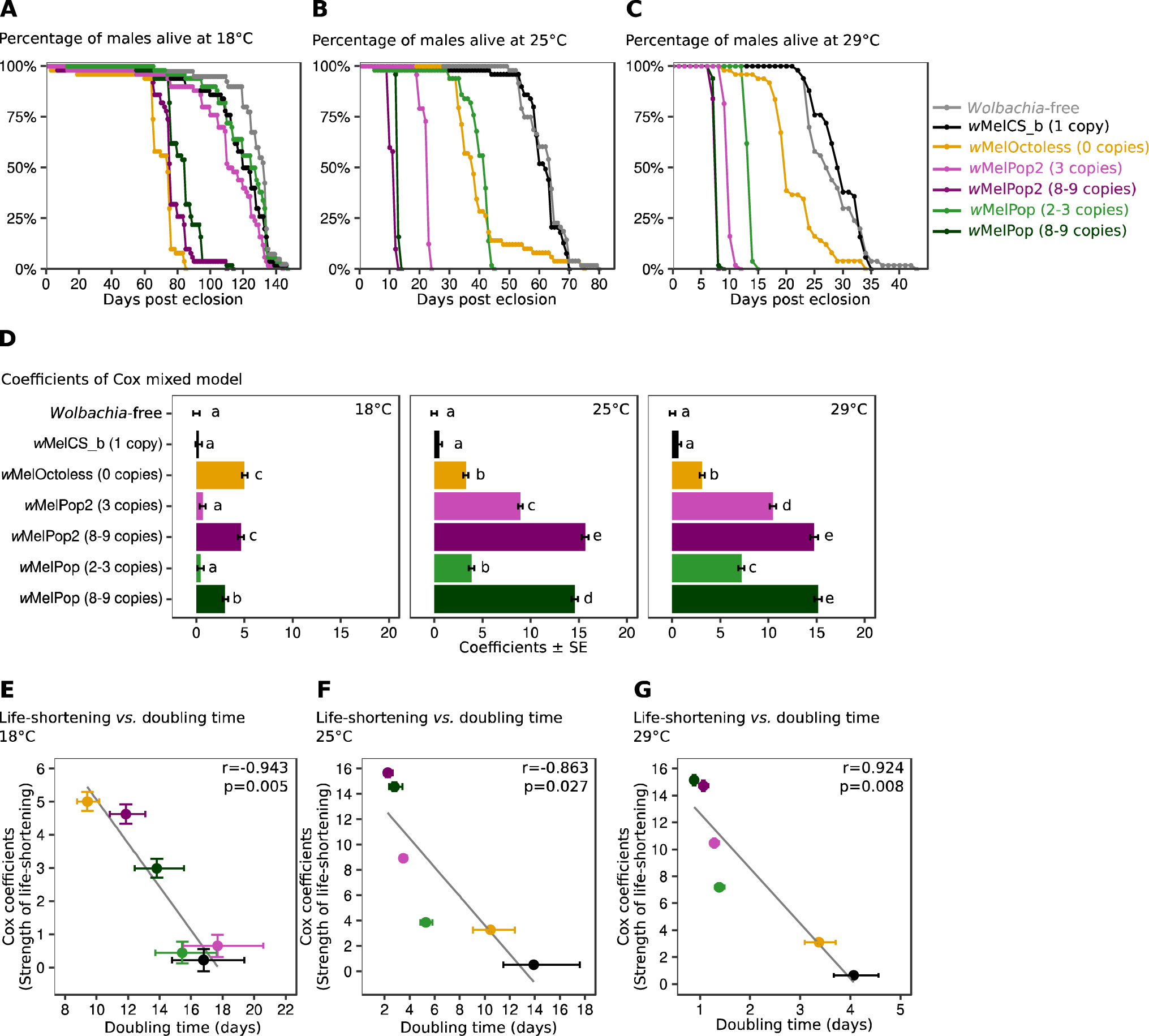
*w*MelOctoless and *w*MelPop2 are pathogenic. Lifespan of males with different *Wolbachia* variants at 18°C (A), 25°C (B), and 29°C (C). For survival analyses, fifty males were collected on the day of eclosion and kept in groups of 10 per vial until all flies died. Flies were transferred to new vials every five days. A full replicate of these experiments is shown in S14A-C Fig. (D) Coefficients of a Cox mixed model, which represent the effect of *Wolbachia* on the lifespan of flies relative to the lifespan of *Wolbachia*-free flies. Both experimental replicates were analysed together. Bars represent the standard error of the coefficient, and letters statistically significant groups after *p*-value correction. (E-G) Correlation between the strength of life-shortening phenotype and *Wolbachia* doubling time at 18°C (E), 25°C (F), and 29°C (G). The y-axis represents the strength of *Wolbachia* life-shortening phenotype (estimated using Cox mixed models) and the x-axis the *Wolbachia* doubling time (in days). The Pearson correlation coefficient (*r*) and its significance (*p*) are given in each panel.

*w*MelOctoless strongly reduces host lifespan at all tested temperatures (*p* < 0.001, each comparison with *w*MelCS_b) (Fig 5A-D, S14 Fig). This deleterious effect is stronger at 18°C, where *w*MelOctoless is the tested variant with the highest impact on lifespan, although very similar and not statistically different from *w*MelPop2 with high Octomom copy number (*p* < 0.001, for all comparisons with other lines, *p* = 1 when compared with *w*MelPop2 with 8-9 Octomom copies). The effect of *w*MelOctoless on host lifespan is weaker at 25°C than at 18°C (*p* = 0.001), and similar at 25°C and 29°C (*p* = 0.95). These results demonstrate that the new over-proliferative *w*MelOctoless also has a cost to the host in terms of lifespan and this effect interacts with temperature, being stronger at lower temperature.

*w*MelPop2, similarly to *w*MelPop, also shortens host lifespan (Fig 5, S14 Fig). The variants containing high copy number of Octomom (8-9 copies) shorten lifespan at all temperatures (*p* < 0.001, for each comparison with *w*MelCS_b). This effect is much stronger at 25°C than at 18°C (*p* < 0.001 for contrasts between both lines and *w*MelCS_b), and similar at 25°C and 29°C (*p* > 0.21 for these contrasts). At these two higher temperatures the lines carrying the variants with high copy number of Octomom have the shortest lifespan of all tested lines (*p* < 0.001 for all comparisons). *w*MelPop2 and *w*MelPop with low copy number of Octomom (2-3 copies) always have a weaker effect on host lifespan shortening than high copy number variants (*p* < 0.001 for all these comparisons). As observed with the high copy number variants, their effect increases with temperature, being stronger at 25°C than at 18°C, and even stronger at 29°C (*p* < 0.05 for these comparisons). In fact, *w*MelPop2 and *w*MelPop with low copy number are only pathogenic at 25°C and 29°C, not at 18°C. These data confirm the association of Octomom region amplification with host lifespan shortening, and the increase in the severity of this phenotype with an increase in Octomom copy number, and an increase in temperature.

In some comparisons *w*MelPop2 and *w*MelPop differ significantly in their pathogenic phenotype (Fig 5D). This could indicate that there were differences in this phenotype between these two lines. Therefore, and as done above in the analysis of proliferation, we repeated this experiment comparing the lifespan phenotype in *w*MelPop2 and *w*MelPop lines with a tightly controlled Octomom copy number (S12 Fig). At 25°C lines both *w*MelPop and *w*MelPop2 with 3 copies of the Octomom region had a shorter lifespan than the line with *w*MelCS_b (*p* < 0.001), and no difference between them (*p* = 0.29). These results show that *w*MelPop2 and *w*MelPop have the same phenotype.

To further demonstrate that the life shortening phenotypes were due to the new *Wolbachia* variants, and not to EMS-induced mutations in the host nuclear genome, we performed reciprocal crosses between flies carrying *w*MelCS_b and flies carrying either *w*MelOctoless or *w*MelPop2 (with 3 or 8-9 copies of Octomom) and followed the survival of their female progeny at 29°C. The female progeny from reciprocal crosses should be identical in the nuclear genome but differ in the *Wolbachia* variant, which is maternally transmitted. The life-shortening phenotype segregated maternally, thus demonstrating that the *Wolbachia* variants carried by the lines are the cause of the phenotypes (S15 Fig). The relative strength of the life-shortening phenotype of the progeny of the reciprocal crosses matches the strength of the phenotypes in the maternal lines, observed in Fig 5 and S14 Fig. Moreover, all the tested lines that inherited *w*MelCS_b had a similar lifespan (*p* > 0.78 for all comparisons), indicating, as expected, no contribution of the host genotype in this set of experiments.

The life shortening phenotype of *w*MelPop has been associated with its over-proliferation and higher titres since its discovery [22]. We tested if these phenotypes were correlated by taking advantage of the data on titres, proliferation and lifespan shortening that we collected from this set of variants at different temperatures. We found a negative correlation between the strength of the life-shortening phenotype and *Wolbachia* doubling time, at all temperatures (Fig 5, |r| > 0.86, p < 0.027, for all correlations). However, we found no significant correlations between the strength of the life-shortening phenotype and *Wolbachia* titres in 0-1 days-old adults (S16 Fig, *p* > 0.05 for all correlations). These results show that over-proliferative variants shorten the host lifespan and the strength of this phenotype correlates with their proliferation rates.

### *Wolbachia* variants with deletion or amplification of the Octomom region provide stronger protection against DCV

Previous studies established a link between *Wolbachia* titres and the strength of anti-viral protection [16–20]. To test if *w*MelOctoless and *w*MelPop2 also provide a stronger protection against viruses, we infected flies with *Drosophila* C virus (DCV), by pricking, and followed their survival for 40 days at 18°C. All *Wolbachia* variants tested provided protection against DCV (CHR, *p* < 0.001 for all comparisons with the *Wolbachia*-free line, Fig 6A-B and S17A Fig), while survival of *Wolbachia*-carrying flies did not differ from control when pricked with buffer, in this time frame (S17B-D Fig, *p* = 0.52 for *Wolbachia* variant effect). *w*MelCS_b was the least protective variant, while *w*MelOctoless was the one providing the highest protection. In general, the over-proliferative *Wolbachi*a variants confer stronger protection to DCV than *w*MelCS_b, although this difference is not always significant (Fig 6B, *p* < 0.001 for all comparisons, except for *w*MelPop (2-3 copies), *p* = 0.11).

**Fig 6.**
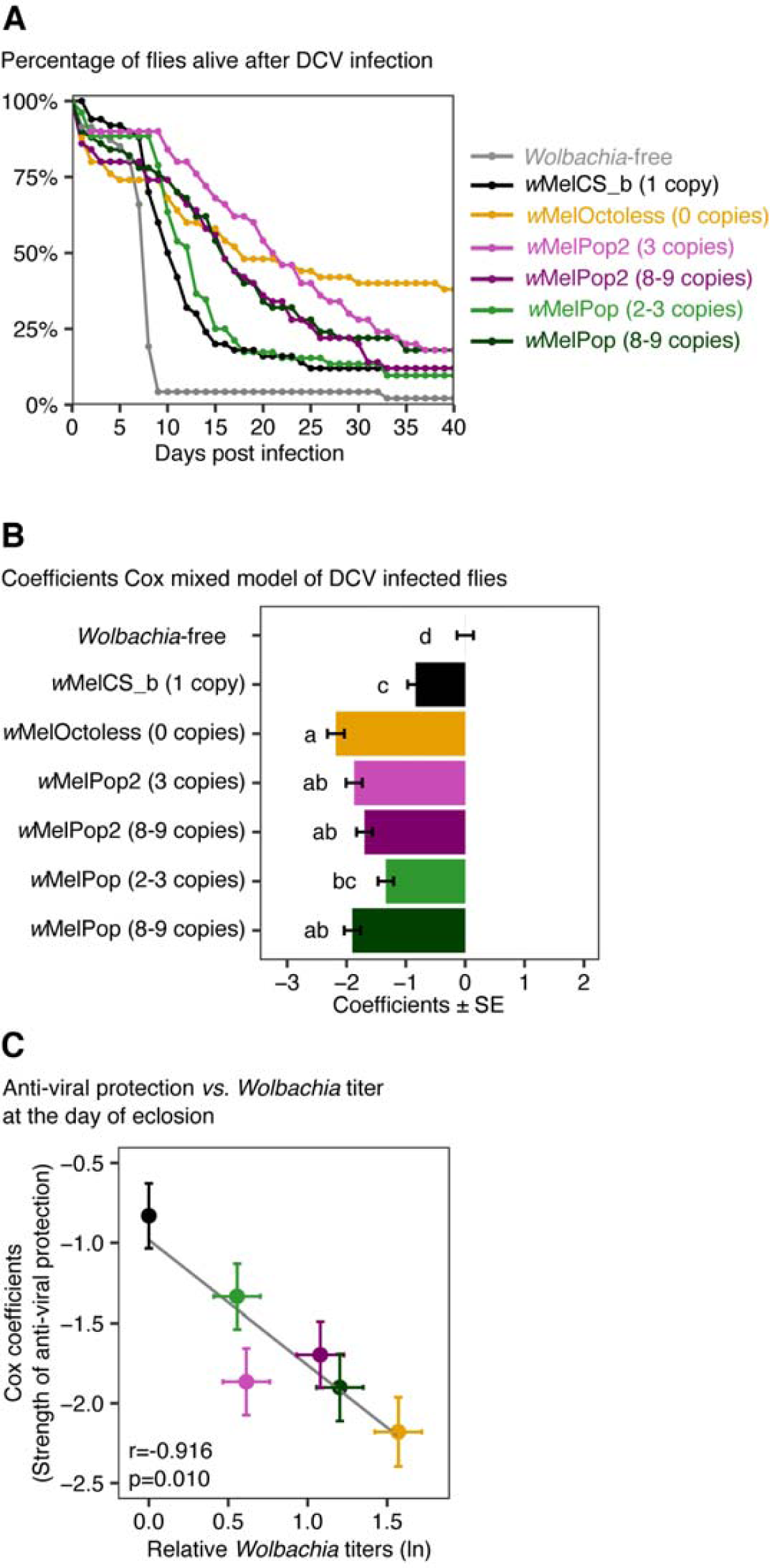
*w*MelOctoless and *w*MelPop2 provide strong protection against DCV. (A) Survival of males carrying different *Wolbachia* variants after a challenge with DCV. Fifty 3-5 days-old *Drosophila* males, per line, were pricked with DCV (10^9^ TCID_50_/ml) and survival curves were determined at 18°C for 40 days. A replicate of the experiment and the buffer-pricked controls are shown in S17 Fig. (C) Coefficients of Cox mixed models of DCV-infected flies. Cox coefficients represent the effect of *Wolbachia* infection on survival relative to the *Wolbachia*-free flies. Bars represent the standard error of the estimate, and the letters the statistical significant groups after *p*-value correction. *Wolbachia* infection improved the survival of DCV-infected flies (p < 0.001). (C) Correlation between the strength of anti-viral protection (represented as the coefficients of a Cox mixed model in the y-axis) and the natural log of *Wolbachia* titres at the day of eclosion, as a proxy for *Wolbachia* titre in the day of infection. The Pearson correlation coefficient (*r*) and its significance (*p*) are given in the panel.

We also tested the correlation between the antiviral protection and *Wolbachia* proliferation, but we found no correlation neither with proliferation estimates at 18°C (*p* = 0.21), the temperature in which the flies were kept after infection, nor at 25°C (*p* = 0.35), the temperature in which flies developed and were kept until being infected with DCV (S16D-E Fig). However, there is a significant correlation between the strength of *Wolbachia*-induced antiviral protection and *Wolbachia* titres in 0-1 days-old flies, a proxy for *Wolbachia* titre at the day of infection, which is 3-5 days-old flies (Fig 6C, *p* = 0.010, |r| = 0.92). Overall, the new over-proliferative variants give more protection to viruses than *w*MelCS_b, and the strength of this protection is correlated with *Wolbachia* levels at the time of infection.

## Discussion

We developed a new forward genetic screen and identified new *Wolbachia* over-proliferative variants. We characterized in detail two of these new mutants, *w*MelPop2 and *w*MelOctoless, and identified the genetic bases of their over-proliferation. *w*MelPop2 had an amplification of the Octomom region, which has been previously shown to lead to over-proliferation in the *w*MelPop variant [16,17]. *w*MelOctoless, on the other hand, had a deletion of this same Octomom region. These results further confirm and develop the complex role of this genomic region in the control of *Wolbachia* proliferation. An extensive phenotypic characterization of two of these lines showed both *Wolbachia* variants to shorten the host lifespan, as well as to increase antiviral protection. Moreover, we show that *Wolbachia* proliferation rate in *D. melanogaster* depends on the interaction between Octomom copy number, the host developmental stage, and temperature. Our analysis also suggests that the *Wolbachia*-induced life shortening and antiviral protection phenotypes are dependent on its rate of proliferation in adults and titres near the time of infection, respectively. These are related, but different, properties of the endosymbiont.

### An unbiased approach for genetically intractable symbionts

Given their dependence on the intracellular niche, *Wolbachia* and most endosymbionts remain non-culturable and genetically intractable, hindering their study. Here we aimed at mutagenizing and screening for new *Wolbachia* variants in the host. We expected that a main difficulty of this approach would be how to identify, via its phenotype, a new mutant present in the *Wolbachia* population within a host. We estimated here that a newly emerged *w*MelCS_b-carrying female harbours approximately 540,000 *Wolbachia* genome copies, which probably corresponds to the same number of *Wolbachia* cells [30]. Since EMS induces random mutations, we expected mosaicism in the *Wolbachia* population at the individual fly level. Each new mutant, when generated, would be a unique cell within these approximately half a million other *Wolbachia* cells. Since the *Wolbachia* phenotypes are normally measured at the individual host levels (e.g. *Wolbachia* titres, antiviral protection), the properties of individual or small numbers of mutant *Wolbachia* could be diluted and unmeasurable.

We hypothesized, however, that over-proliferating *Wolbachia* cells could overtake the population and that the resulting higher titres could be detectable. Indeed, fast proliferative *Wolbachia* can be selected at the level of a single host [31]. To increase the probability of isolating rare over-proliferating *Wolbachia* variants, we also relied on the bottleneck imposed in the vertical transmission of *Wolbachia*. We calculated here that single embryos carry approximately 3,000 *Wolbachia* genomes, which is consistent with previous estimates [29]. We also treated flies with tetracycline before mutagenesis to further reduce the *Wolbachia* population. We expected this additional endosymbiont titre reduction to enhance genetic drift and potentially enrich or lead to the fixation of rare *Wolbachia* variants. By screening at the immediate progeny (F1) of EMS-treated females or three generations later (F4) we were able to select new over-proliferative *Wolbachia* mutants.

### Genetic bases of *Wolbachia* over-proliferation

After discarding the possibility that mutations in the host nuclear or mitochondrial genomes were the cause of *Wolbachia* over-proliferation, we performed *de novo* assembly of the ancestral, *w*MelCS_b, and the new mutant variants, *w*MelPop2 and *w*MelOctoless. *w*Mel and *w*MelPop were also assembled. The assemblies generated complete full chromosomes of these *Wolbachia* and allowed us to identify single nucleotide differences and structural differences between these genomes. To validate our genome assembly pipeline we compared our *w*Mel genome assembly to the reference *w*Mel genome. We identified only seven indels and two SNPs, which we confirmed to be present in our line, by Sanger sequencing. Our assembly results also showed two previously identified SNPs between *w*MelCS_b (but also the new *w*MelCS_b derived variants *w*MelOctoless and *w*MelPop2) and *w*MelPop. Additionally, our assembly provides an improvement over the previous *w*MelPop genome [25].

The only differences between the new over-proliferative variants and *w*MelCS_b were structural differences in the Octomom region. *w*MelPop2 has an amplification of this region. The assembly confirms that the Octomom region is amplified in tandem [17], and that all copies are located in the *Wolbachia* genome. Previously we showed that amplification of the Octomom region and the degree of this amplification determined *w*MelPop over-proliferative phenotype [16,17]. Moreover, reversion of *w*MelPop Octomom copy number to one, through selection, resulted in loss of both over-proliferation and cost to the host, making the variant phenotypically identical to *w*MelCS_b, which also carries a single copy of Octomom [17]. We now show that Octomom amplification in *w*MelCS_b also leads to an over-proliferative phenotype. Moreover, *w*MelPop and *w*MelPop2 variants, carrying the same copy number of Octomom, have identical phenotypes. These findings are unsurprising since *w*MelCS_b, the ancestral of *w*MelPop2, and *w*MelPop share an almost identical genome, differencing on two synonymous SNPs and Octomom amplification. Hence, our results further confirm the role of amplification of Octomom region in the over-proliferation of *Wolbachia*.

*w*MelOctoless, on the other hand, has a deletion of the entire Octomom region. The deletion leaves behind one of the direct repeats flanking the Octomom region [16], suggesting that excision might have been mediated by recombination. The causal link between Octomom deletion and over-proliferation is further supported by an additional independent over-proliferative variant isolated in the screen, *w*MelOctoless2. This *Wolbachia* variant also has a deletion of Octomom as the only difference with *w*MelCS_b. These data show that deletion of the Octomom region also leads to an over-proliferative phenotype in *Wolbachia*. Thus, we identified the second known *Wolbachia* mutation with a clear link between genotype and phenotype.

The mutations identified in the new variants are deletions and amplifications. We did not detect any new SNPs in these variants, even if EMS is expected to mainly induce single nucleotide mutations [32]. Thus, it is possible that the deletion or amplification of the Octomom region in these over-proliferative variants were independent of the EMS treatment. For instance, loss of the Octomom region has been twice reported, in cell culture [25,33], suggesting that it may occur spontaneously. Yet, chemical mutagens such as EMS can activate DNA damage response and transposable elements [34], and some genes of the Octomom region and its flaking genes are predicted to be potentially involved in transposition and DNA repair [16]. Therefore, we cannot rule out that the EMS treatment induced the mutations in this genomic region.

### Opposing mutations lead to a similar *Wolbachia* over-proliferative phenotype

Both the deletion or the amplification of the Octomom region causing an over-proliferative phenotype seems to be a paradox. The resolution of this paradox and the mechanisms leading to these phenotypes will rely on the functional characterization of genes in the Octomom region. These genes may be involved in interaction with the host, transcriptional regulation or DNA repair [16].

One possibility is that the amplification of the Octomom region and over-expression of a particular set of genes in this region lead, mechanistically, to the same result as the absence of the genes. There are many examples of over-expression of a gene leading to a dominant negative phenotype. For instance, both over-expression or loss of a protein forming a gradient, abolish the gradient [35]. Also, over-expression of a member of protein complex may lead to loss of stoichiometry and therefore loss of functional complexes [36]. Another possibility is that the loss and amplification of different genes in the Octomom region lead to the over-proliferative phenotype. The second hypothesis is supported by the fact that *w*MelPop2 and *w*MelOctoless have similar but not identical phenotypes, and interact differently with temperature. Furthermore, *w*MelPop and *w*MelPop2 have a higher rate of proliferation than *w*MelOctoless at 25°C and 29°C. Therefore, at these temperatures, the phenotype of Octomom amplification is stronger than the phenotype associated with the Octomom region loss-of-function.

Although complex, these results help to explain some of the data from an over-proliferative variant trans-infected in *Aedes aegypti*. The *D. melanogaster* variant *w*MelPop was transinfected into *Aedes albopictus* cells, and then transinfected into *A. aegypti*. In the process of cell culture adaptation the Octomom region, which was amplified in *w*MelPop, was deleted [25]. The *w*MelPop-PGYP variant in *A. aegypti* lacks, therefore, the Octomom region. Nonetheless, this variant still over-proliferates and is highly pathogenic in *A. aegypti* [21]. If one assumed the same genetic basis for the pathogenicity of *w*MelPop in *D. melanogaster* and *w*MelPop-PGYP in *A. aegypti*, one might conclude that the Octomom region was not related with these phenotypes [25]. However, similarly to *w*MelPop/wMelPop-PGYP variants our results with *w*MelCS_b and the mutant variants show that both amplification and deletion of the Octomom region lead to increased *Wolbachia* pathogenicity. Thus, deletion of the Octomom region in *w*MelPop-PGYP may explain why this variant is also pathogenic. However, *w*MelPop-PGYP also accumulated other mutations when passaged in cell culture, and has many other genetic differences with *w*Mel [25]. These may also contribute to *w*MelPop-PGYP being more pathogenic than *w*Mel in *A. aegypti*.

The Octomom region has the properties of a genomic island: it is not part of the *Wolbachia* core genome, since many *Wolbachia* strains lack this region, and seems to be horizontally transferred between *Wolbachia* strains [16,37,38]. Although the *w*Mel strain can lose this region and remain viable in laboratory conditions (here and [25,33]), natural variants of *w*Mel without Octomom are not known [39]. Its over-proliferative phenotype, and the associated shortening of host lifespan, may lead to a fitness disadvantage to a host harbouring such mutant. This in turn may lead to loss of these variants from the host natural populations. Therefore, this genomic region, absent in many other strains of *Wolbachia*, became addictive to the *w*Mel strain through its integration in the regulation of *Wolbachia* proliferation.

### *Wolbachia* proliferation, pathogenicity, and antiviral protection

We found a complex interaction between temperature and proliferation of the different *Wolbachia* variants in adults. *w*MelOctoless proliferates faster than *w*MelCS_b, to a similar extent, at all temperatures. *w*MelPop and *w*MelPop2, however, strongly interact with temperature and proliferate much faster at higher temperatures. We also confirmed here that the degree of Octomom amplification in these variants modulates the proliferation rate [17], and interacts with temperature.

Throughout the range of tested temperatures, the different variants have very different proliferation rates in adults. At 25°C, where we observed the highest variation, the titres of *w*MelPop2 double every 2 days, while *w*MelCS_b titres double every 14 days. In contrast, during larval development the proliferation rates of *w*MelCS_b, *w*MelOctoless, and *w*MelPop2, are very similar and much faster than in adults. During development, at 25°C, *Wolbachia* titres double every 12 to 16 hours. These results show that *w*MelCS_b, which has a relatively low proliferation rate in adult flies, is capable of very fast proliferation during larval development, when the host is rapidly growing. In *Brugia malayi*, *Wolbachia* titres also increase rapidly during the first few weeks of the nematode’s development [30]. Thus, rapid proliferation during immature stages may be a conserved *Wolbachia* strategy to recover from the bottleneck imposed during maternal transmission. This observed coordination between *w*MelCS_b proliferation and *D. melanogaster* developmental stage may be due to *Wolbachia* directly responding to host developmental cues, to differences in the metabolic profile of larvae and adults, or to differences in host cell divisions rates. *Wolbachia* could also control its proliferation in response to its own population density within the host.

While wMelPop and wMelPop2 proliferate similarly to *w*MelCS_b during larval stages, they over-proliferate in adults. Thus, this over-proliferative phenotype of variants with amplification of the Octomom region can be interpreted as an inability to properly respond to the conditions of the host adult stage. On the other hand, the deletion of Octomom seems to lead to a similar increase in the proliferation rate during development and adult life, although the difference with *w*MelCS_b is not significant during development in our analysis.

The new over-proliferative variants shorten the lifespan of *D. melanogaster*, as *w*MelPop does. Furthermore, we showed this phenotype to result from the interaction of *Wolbachia* genotype and temperature. *w*MelOctoless had a similar life shortening phenotype at all temperatures, although it was stronger at 18°C. *w*MelPop and *w*MelPop2 responded strongly to temperature, being much more costly at higher temperatures, as shown before for *w*MelPop [17,22,40]. The extent of Octomom amplification also influenced this interaction. While low copy number variants had no phenotype at 18°C, the high copy number *w*MelPop and *w*MelPop2 are pathogenic also at this temperature. Therefore these variants can also be pathogenic at 18°C, contrary to previous data [28,40,41]. Interestingly, we find at all temperatures a significant correlation between the proliferation rate of the *Wolbachia* variants and the life shortening phenotype. The faster the variants proliferate the shorter the host lifespan.

All the over-proliferative variants also increased antiviral resistance, with *w*MelOctoless conferring the strongest protection. This phenotype correlated poorly with proliferation rates at 25°C or 18°C, the temperature before and after infection with DCV, respectively. However, the strength of the antiviral resistance correlated with the titres of *Wolbachia* near the day of infection. Thus, the cost of harbouring *Wolbachia* in terms of lifespan, and the benefit of the antiviral protection, correlate with related but different parameters. In the future it will be important to understand why these different correlations. For instance, *Wolbachia* titres at the point of viral infection are probably important because the anti-viral protection is observed early in the infection [42]. *Wolbachia* proliferation rate could impact longevity either by the cumulative cost of the proliferation process itself or by determining the time to reach a lethal threshold of *Wolbachia* titres. Nonetheless, these results indicate that it may be possible to select for highly protective *Wolbachia* variants without necessarily having a high cost to the host. These would be *Wolbachia* variants with high titres but low proliferation in adults. Such variants would be particularly useful in the use of *Wolbachia*-transinfected mosquito to prevent arboviruses transmission.

In summary, our results show the feasibility of forward genetic screens to study *Wolbachia* biology. Similar strategies may be used in the future to study other aspects of *Wolbachia*-host interactions or the biology of other genetically intractable endosymbionts. The new over-proliferative variant *w*MelPop2 confirms the causal link between amplification of the Octomom region and *Wolbachia* over-proliferation. Whereas the new loss-of-function mutant *w*MelOctoless reveals that this region is also required to control *Wolbachia* proliferation. These results give new insight on the complex role this genomic region plays in *Wolbachia* biology. Moreover, this collection of variants, similarly to an allelic series, allow a finer dissection of the consequences of *Wolbachia* over-proliferation to the host.

## Materials and Methods

### Fly genotypes, infection status, and maintenance

Flies were reared on fly food, supplemented with live yeast, at 25°C, 70% humidity. Fly food was composed of molasses (37.5g/L), sugar (62.5g/L), cornflour (58.3g/L), yeast extract (16.7g/L), and agar (8.3g/L) in distilled water. The mixture was sterilized by autoclaving and cooled to 45°C. For each litre of food, we added 29.2 mL of a solution with 100g of methylparaben and 0.2g of Carbendazim for 1L absolute ethanol.

All fly stocks used had the Drosdel *w*^1118^ isogenic background [16,43]. The bacterial community associated with the fly stocks was homogenized as in Pais *et al.* [44], with minor modifications. Briefly, we collected eggs for 6 hours in fresh agar plates with live yeast and sterilized the eggs surface by consecutive washes on 2.1% sodium hypochlorite (NaOCl) solution (10 minutes), 70% ethanol (5 minutes) and sterile water (5 minutes). Next, we transferred axenic eggs to sterile fly food supplemented with 40μL of 1:1 overnight culture of *Acectobater* OTU 2753 and *Lactobacillus* OTU 1865 [44]. We confirmed the presence of these bacterial species by squashing five females aged 3–6 days in sterile 1x PBS, plating 30μL of the lysate in mannitol plates, incubate them at 25°C for 72h, and identify bacteria by colony morphology.

### Selection of *D. melanogaster* lines carrying *Wolbachia* with specific Octomom copy number

To select for flies carrying *w*MelPop and *w*MelPop2 with a desired Octomom copy number, we proceeded as in Chrostek and Teixeira [17], with minor modifications. Briefly, we allowed 5–20 virgin females to cross with 2–3 *Wolbachia*-uninfected males of the Drosdel *w*^1118^ isogenic background in individual vials, and lay eggs for 3-4 days. Females were then collected in individual tubes for DNA extraction and Octomom copy number determination by qPCR. The progeny of females with specific copy numbers were then followed-up.

### Determination of time for *Wolbachia* titres recovery

Flies with *w*MelCS_b developed in fly food supplemented with tetracycline at the concentrations 1.5625µg/ml, 3.125µg/ml, 6.25µg/ml, 12.5µg/ml, 25µg/ml, and 50µg/ml. Three isofemale lines were established from each dose. In the F1, we randomly selected four virgin females for egg-laying and *Wolbachia* titre measurement using qPCR. From this moment on, the flies were kept on fly food without tetracycline. We set up the next four generations using the progeny of a female with the median *Wolbachia* titres.

### Forward genetic screen

We attempted to mutagenize *Wolbachia in vivo* by feeding its host with the mutagen EMS. DrosDel *w*^1118^ isogenic flies carrying *w*MelCS_b were raised in standard fly food or fly food supplemented with tetracycline (from 1.5625 to 12.5µg/ml). Virgin females were collected, starved for 6h, and then fed EMS concentrations ranging from 10 to 8,000mM diluted in 1% sucrose. Control flies fed on sucrose solution only. A dye was added to the feeding solution to confirm intake and feeding proceeded for 13h (overnight).

EMS-fed females (G0), and control females, were mated individually with 2–3 *Wolbachi*a-free Drosdel *w*^1118^ isogenic males, egg-laying was allowed for 3–4 days, and parents discarded. From the F1 progeny, we collected virgin females, mated them individually with 2–3 *Wolbachi*a-free Drosdel *w*^1118^ isogenic males, and allowed egg laying for 3-4 days. These females were collected when 10 days old, and *Wolbachia* titres determined by qPCR. We followed the progeny of F1 females showing 50% or higher increase in *Wolbachia* titres relative to control flies in the same conditions. We have also transferred the progeny of these F1 for three more generations, without selection, and repeated the determination of *Wolbachia* titres in F4 females. In the same batch of experiments we may have tested more than one F1 or F4 progeny from each G0 female. Hence, over-proliferative *Wolbachia* variants isolated in the same batch of treated females may be a result of a single event in the G0 female.

### Real time quantitative PCR

DNA extraction for qPCR was performed as described before [17].

The qPCR reactions were performed in the QuantStudioTM 7 Flex (Applied Biosystems). The reaction mix and PCR program used were described before [16]. The specificity of the amplicons was confirmed by analysing the melting curve profile.

Relative levels of the target genes was determined using the Pfaffl method [45]. To quantify relative *Wolbachia* titres we used *Drosophila RpL32* gene as calibrator, and *Wolbachia wsp* as the target gene. To determine the copy number of the Octomom region, *Wolbachia wsp* gene was used as the calibrator and *WD0513* used as the target gene. For determination of copy number of other Octomom region genes, or control genes, *wsp* was also used as a calibrator.

For absolute quantification of *Wolbachia* genome copies the full-length *Wolbachia wsp* gene was cloned into a pMT/V5 *Drosophila* expression vector (Invitrogen). The plasmid was amplified in *Escherichia coli* strain DH5-α, purified using midiprep (QIAGEN) and its concentration determined using Qubit® 2.0 (Thermo Fisher Scientific). Molecular weight of the plasmid was calculated assuming a nucleotide average weight of 325 Da to determine the number of plasmid molecules in the calibration curve. Standard curves of 1:10 serial dilutions were run to calibrate the assay each time.

Primers used in qPCR reactions are given in S8 Table.

### Determination of *Drosophila* lifespan

For each replicate, a total of 50 males or 50 females were collected on the day of eclosion. Flies (10 per vials) were then incubated at 18°C, 25°C or 29°C, and transferred to new fresh vials every four (females) or five (males) days. The number of dead flies was recorded daily until all the flies died. Censored observations (i.e. flies lost or trapped in the vial plug) were recorded and taken into account during data analysis.

### Protection against *Drosophila* C Virus

We produced and titrated the *Drosophila* C virus solution as described in Teixeira *et al.* [8]. We infected 50 3-5 days old males by dipping insect needles (Austerlitz Insect Pins) into a virus solution (10^9^ TCID_50_/ml in 50mM Tris-HCl, pH 7.5) and pricking flies anaesthetized under CO_2_ in the thorax. An equal number of males were pricked with a buffer only solution (50mM Tris-HCl, pH 7.5) and served as controls. After pricking, flies were incubated in groups of 10 individuals per vial, and kept at 18°C. Survival was followed as above.

### *Wolbachia* proliferation during development and in adults

To determine *Wolbachia* growth during development, flies carrying *w*MelCS_b, *w*MelOctoless, and *w*MelPop2 (high-copy), laid eggs for 2 hours in apple juice agar plates supplemented with live yeast. Eggs were transferred to fly food-containing bottles and allowed to develop at 25°C. For *Wolbachia* titre assessment, we sampled eggs (2 hours), L1 larvae (24 hours later), newly moulted L2 larvae (48 hours later), L3 larvae (72 and 84 hours), white prepupae pupae (120 hours), P8 staged pupae (168 hours), and newly eclosed adult males and females (240 hours). Ten samples per time point were analysed. Samples included ten individuals each for eggs and larvae and one individual each for pupae and adults. To collect newly molted larva, all larvae of the target stage were discarded at the respective time-point and the newly molted larvae were collected within two hours after the established time-point. The white prepupae were collected by staging and the remaining stages were collected according to the set time-point. Except for adults, which were collected within 24 hours post-emergence, all samples were collected within two hours interval.

For assessment of titres dynamics in adult flies, newly eclosed males, raised at 25°C, were incubated at 18°C, 25°C, and 29°C. Flies were collected every three (29°C), seven (25°C) or ten days (18°C) for *Wolbachia* titre measurement. Ten individuals were processed for each time point, and the experiment was performed twice. Each sample consisted of a single fly.

### *Wolbachia* genomes sequencing and quality control

For *Wolbachia* genomic sequencing (Illumina and Oxford Nanopore), we enriched the sample for *Wolbachia* cells before DNA purification. To this end, approximately 500 10-days old flies were squashed for 5 minutes in 10ml Schneider’s Insect Medium (Thermo Fisher Scientific) using glass beads. Next, we pelleted host debris by centrifugation at 1,000g for 5 min and filtered the supernatant solution thought a 5µm pore. Bacterial cells were pelleted by centrifugation at 13,000rpm for 15 minutes, and DNA was extracted. All centrifugations were carried out at 4°C. DNA was extracted with a phenol-chloroform isolation protocol and resuspended in 10mM Tris-HCl (pH 8).

Sequencing was performed at the Genomics Facility at the Instituto Gulbenkian de Ciência, Portugal. Both Illumina and Oxford Nanopore sequencing was done on genomic DNA extracted from the same biological material. Illumina 300bp paired-end libraries were prepared using the Pico Nextera kit according to the manufacturer’s instructions and sequenced with MiSeq. Data quality was assessed via FastQC v.0.11.5 [46] and reads were trimmed using Trimmomatic v.0.36 [47]. Genomic samples for Oxford Nanopore sequencing were processed with minimal shearing to maximize the size of fragments in the libraries. After ligation (kit SQK-LSK108), libraries were sequenced in MinION Mk1b portable sequencing device using SpotON flow cell (R9.4.1). The status of the sequencing pores was monitored using MinKNOW (v2.0.1). Sequencing lasted for up to 48 hours. Albacore (v2.3.1) and Porechop (v0.2.2) were used for base-calling and read trimming, respectively.

### Genome assemblies and comparison

Illumina and Oxford Nanopore reads were first mapped to *D. melanogaster* genome (BioProject: PRJNA13812) using BWA mem v0.7.12-r1039 [48] and minimap2 v2.17-r941 [49], respectively. Reads mapping *D. melanogaster* genome were removed from the datasets before proceeding with *Wolbachia* genome assembly. We used Unicycler v0.4.8-beta [50] assembly pipeline on the remaining reads in order to assemble the *Wolbachia* genomes. Briefly, Unicycler uses Illumina reads to produce a repeats-limited image graph using Spades v3.9.0 [51], which was further refined through Bandage v0.8.1 [52]. Both small short nodes and nodes with no homology with *w*Mel genome (AE017196.1) upon blastn v2.8.1+ [53] search were removed. Next, repeats were resolved by bridging Spades assemblies with Oxford Nanopore long reads. The resulting draft assemblies were polished using Racon v1.3.1 [54] and Pilon v1.23 [55] and rotated so that genomes begin at the *dna*A gene (draft 1 genomes, not published).

We further refined our genome assemblies by mapping the Illumina reads to the corresponded draft genomes to identify mismatches, which were later corrected via Sanger sequencing (draft 2 genomes, not published). Primers used are in S9 Table. Next, we compared the draft 2 genome assemblies by aligning *w*MelPop, *w*MelCS_b, *w*MelPop2, and *w*MelOctoless using Mauve v2.4.0 [56]. The differences between these genomes could correspond to differences between *Wolbachia* variants or still genome assembly artefacts. All detected differences were analysed by Sanger sequencing (primers in S9 Table). There were no confirmed SNPs or small indels between *w*MelCS_b, *w*MelPop2, and *w*MelOctoless. However, we identified and confirmed using Sanger sequencing two predicted SNPs between *w*MelPop and the other *Wolbachia* variants. The genomes were corrected with the Sanger sequencing information and published (BioProject: PRJNA587443).

We further tried to identify mutations in the over-proliferative variants following a previously published pipeline [57]. It consisted of mapping quality checked reads to a reference genome using BWA mem v0.7.12-r1039 algorithm [48] and saving the output as Sequence Alignment/Map file format (SAM). After conversion to the Binary Alignment/Map format (BAM), the file was sorted, duplicates removed and indexed using SAMtools v0.1.19 [58]. Next, we generate mpileup files, also using SAMtools (option ‘-d 1,000,000’), after which was converted to Variant Call Format (VCF) files using BCFtools v1.9-209. We visually confirmed all inferred mutations in IGV v2.4.2 [59]. We did not consider mutations associated with homopolymer regions or in regions with low coverage (<10X). The set of mutations were compared between *Wolbachia* variants using custom Python and R scripts. The only difference we detected between these genomes was higher coverage or deletion of Octomom region.

To compare the set of mutations in the mitochondria of flies infected with different *Wolbachia* variant, we mapped Illumina reads to the *D. melanogaster* Release 6 genome sequence (KJ947872.2:1–14,000) and proceed as previously. Mutations following the criteria previously described were also compared by using custom scripts.

### Statistical analysis

All the statistical comparisons were performed in R v4.0.0 [60].

To compare *Wolbachia* titres across multiple groups, we used linear models (LM) or linear mixed models (LMM). The effect of EMS on *Wolbachia* titre was tested using non-linear regression. We estimated the doubling time of *Wolbachia* variants using the equation log(2)/β, with β being the coefficients of an exponential model.

The lifespan datasets and survival curves after challenge with DCV were analysed with mixed effect Cox models [61].

The significance of correlations were tested using Pearson’s correlation coefficient.

If multiple comparisons were necessary, the p-values were adjusted as proposed by Holm [62]. When multivariate techniques were applied, all the relevant covariates were included in the model, and the final model was selected as proposed by Burnham & Anderson [63].

All statistical analysis and supporting data is deposited in https://doi.org/10.6084/m9.figshare.14079920.v1 [64].

## Acknowledgments

We are thankful to the Bioinformatics Unit and the Genomics Facility at the Instituto Gulbenkian de Ciência for the support in the sequencing and assembly of the *Wolbachia* genomes. We are thankful to the Fly Facility at Instituto Gulbenkian de Ciência for support to the *Drosophila* work.

## Supporting information

**S1 Fig.**
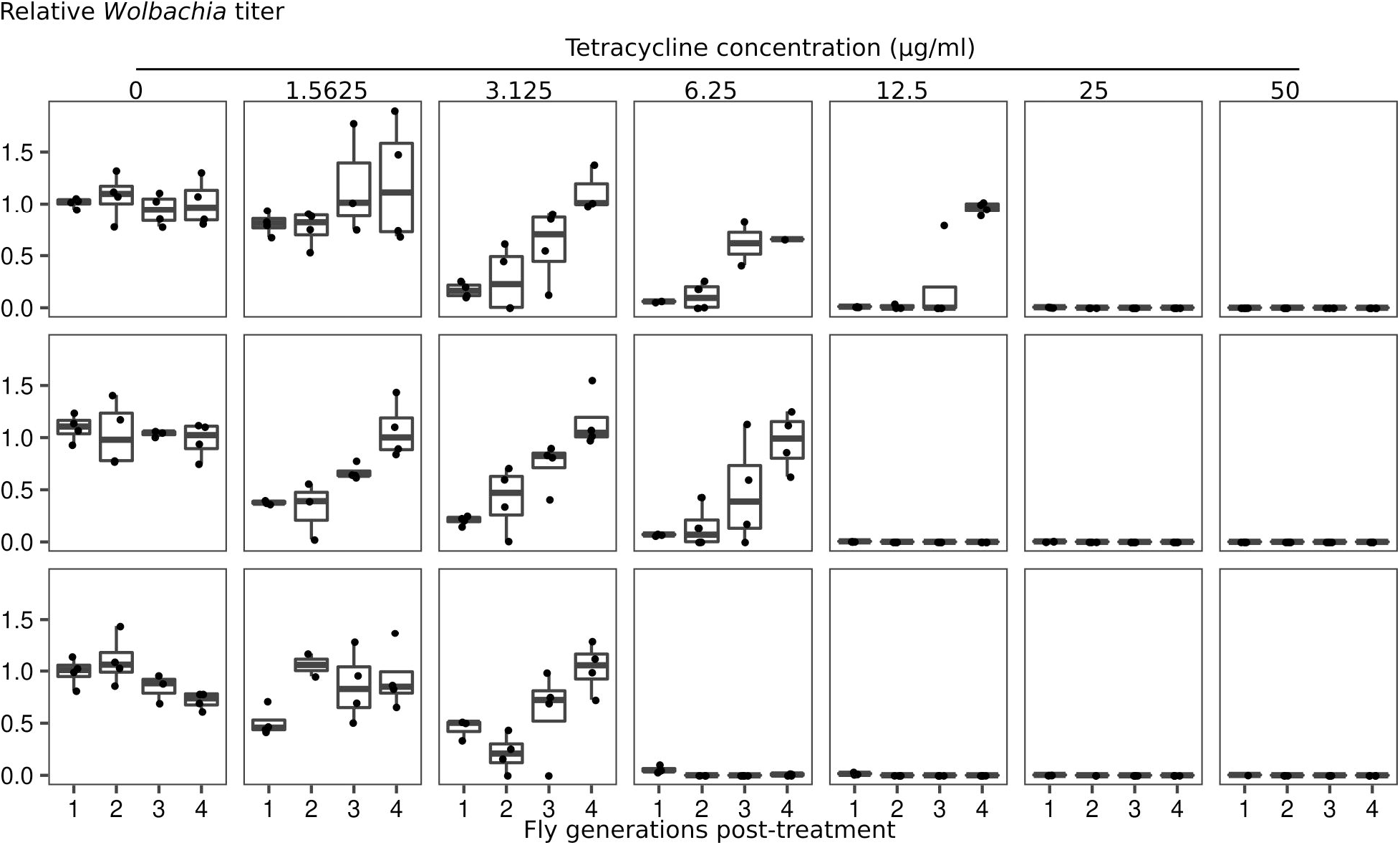
*Wolbachia* recovers from severe titre reduction within four fly generations. Relative *Wolbachia* titres of the progeny of tetracycline-treated flies. *w*MelCS_b-carrying females laid eggs in food containing varying doses of tetracycline. The progeny of three females were used to set up the experiment. At the first generation, four females were randomly selected for egg-laying in antibiotic-free fly food and *Wolbachia* titre was measured using qPCR. Titres of untreated females were used to normalize the qPCR results. The progeny of a female with the median titre was used to set up the next generation. *Wolbachia* titre in the F1 was significantly determined by the concentration of the antibiotic (*p* < 0.001 for all doses compared with control at generation 1), but recovered to normal within four fly generations (*p* > 0.05 for all doses compared with control at generation 4).

**S2 Fig.**
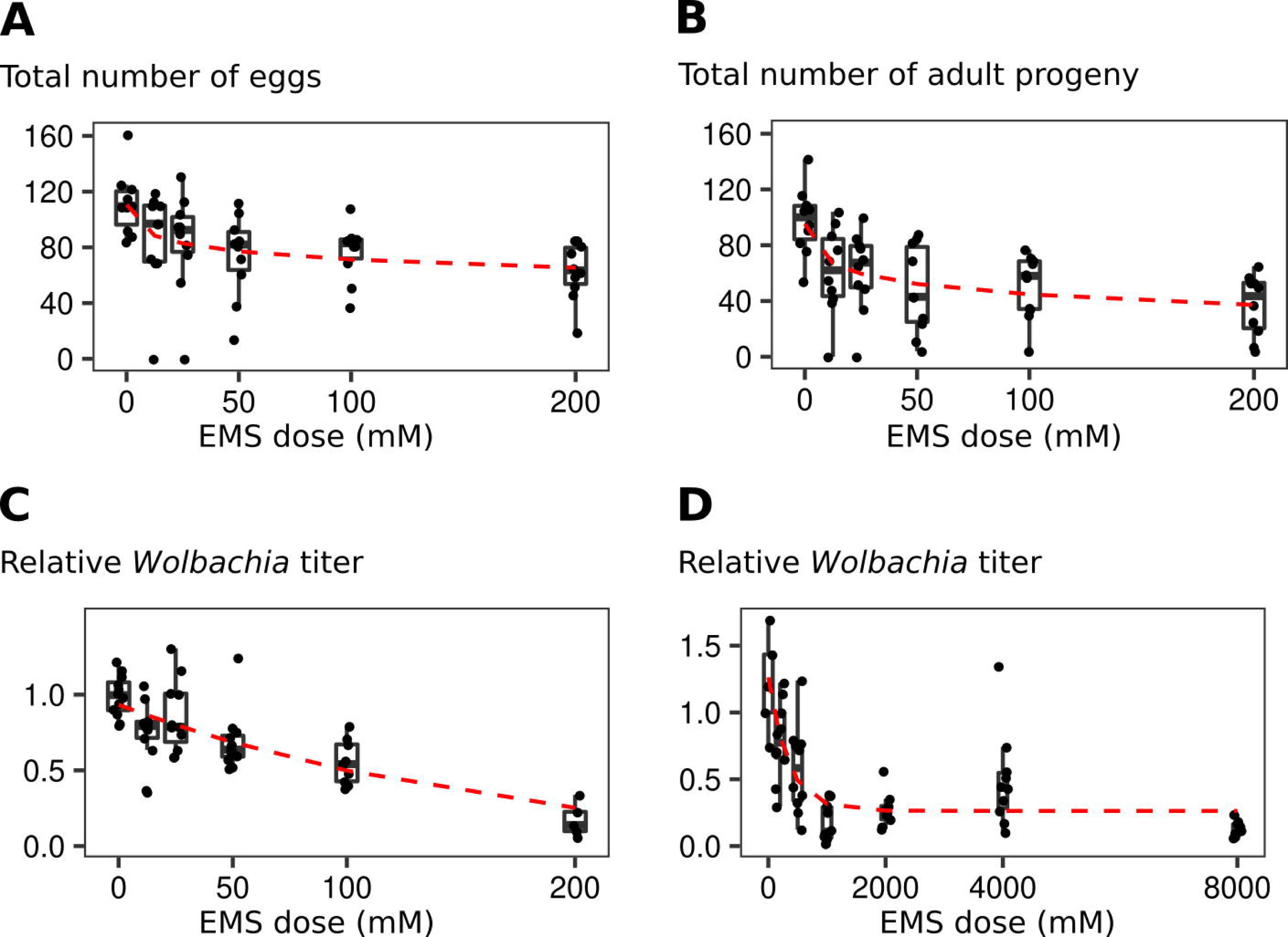
EMS decreases female fecundity and *Wolbachia* titre in the next generation in a dose-dependent manner. The total number of eggs (A) and adults (B) from females treated with varying doses of EMS. The reproductive output of 10 females was determined in the first ten days after EMS treatment by daily transferring females to new vials for egg laying. Females fed on a sucrose solution served as controls. Each dot represents the total number of eggs (A) or adults (B) laid by individual females during ten days. The effect of EMS on the reproductive output of females was estimated using a non-linear model and was highly significant (p < 0.001 for both numbers of eggs and adults per female). (C and D) *Wolbachia* titres in the F1 progeny of females treated with varying EMS doses. *Wolbachia* titre was quantified on individual females (n = 5–13 per dose), after laying eggs for three days. *Wolbachia* titres were normalized against the titres of untreated females. Dashed red lines represent the mean value predicted using non-linear models. The effect of EMS on *Wolbachia* titres in the next generation was highly significant (*p* < 0.001 for both panels).

**S3 Fig.**
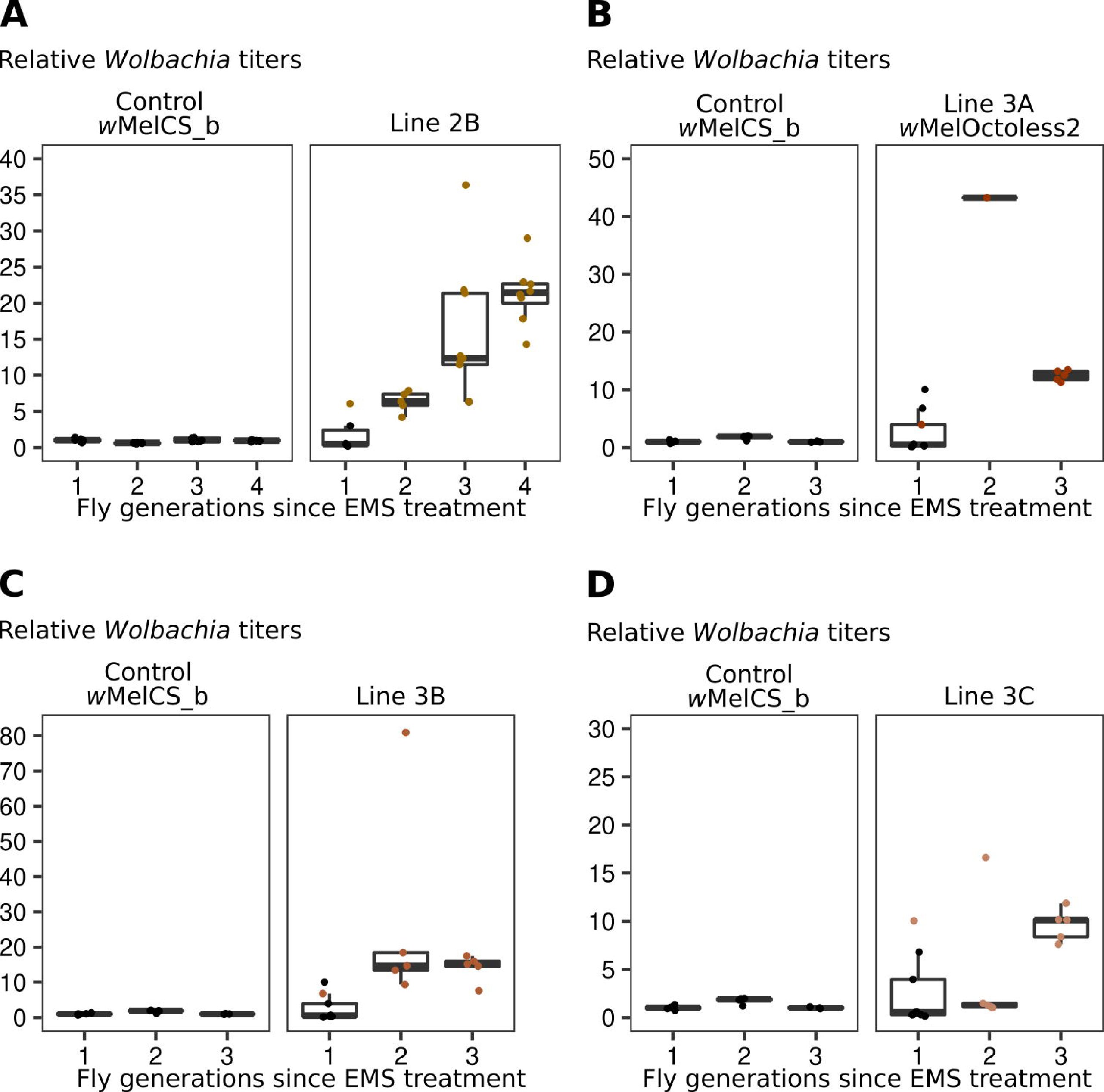
Isolation of over-proliferative *Wolbachia* variants. (A-D) Relative *Wolbachia* titres in a control (wMelCS_b) and EMS-treated *D. melanogaster* lines. Flies to set up the next generation was selected as described for Fig 1. Line 2B was isolated in the same batch as Line 2A (wMelOctoless) and they may be not independent. Likewise, Lines 3A (wMelOctoless2), 3B, and 3C were also isolated in a same batch.

**S4 Fig.**
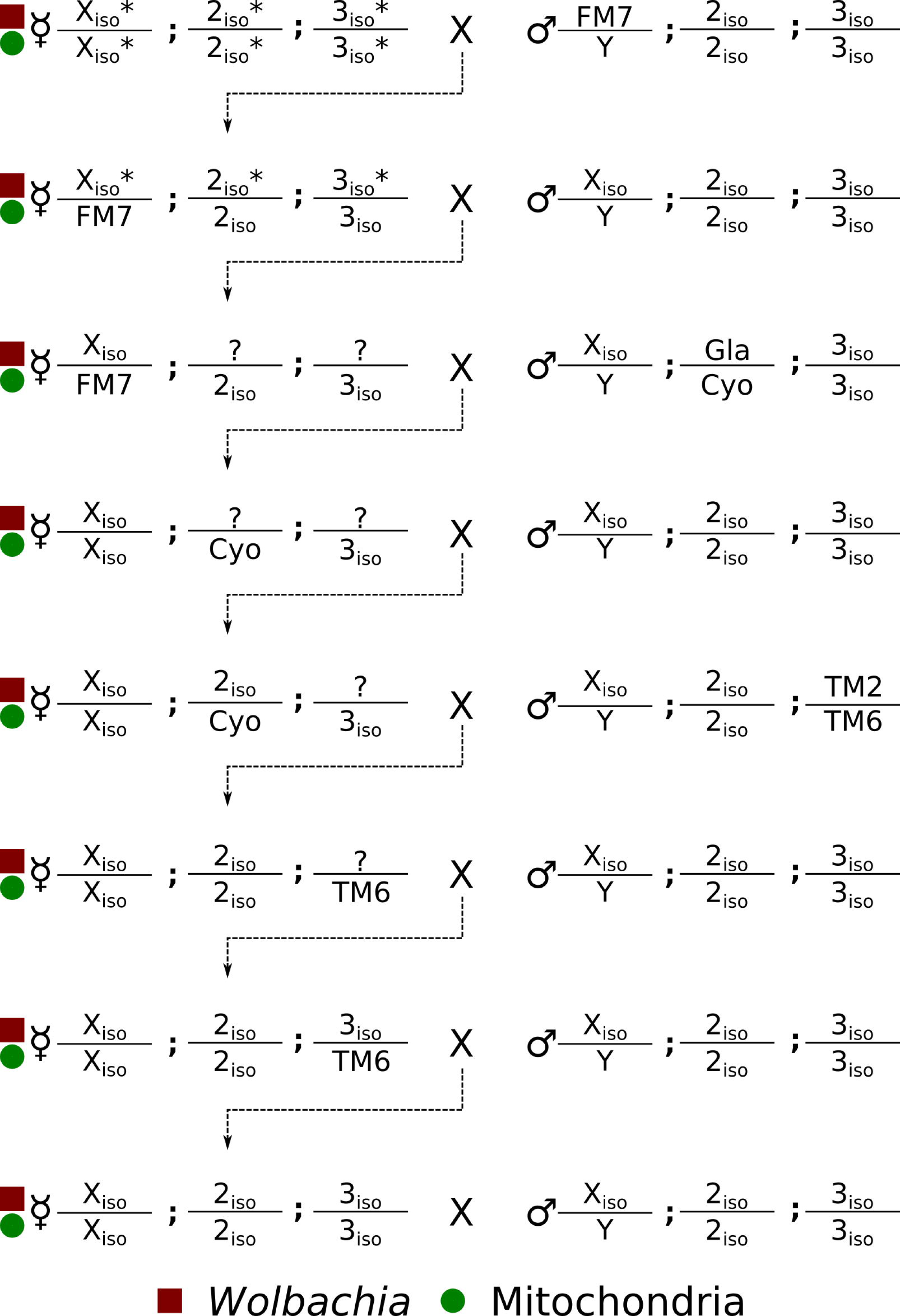
Generation of isogenic *D. melanogaster* lines with *w*MelPop2 and *w*MelOctoless. The first, second and third chromosomes of flies carrying *w*MelPop2, *w*MelOctoless, and *w*MelOctoless2 were replaced through the use of balancer chromosomes. *Wolbachia* infection (and also mitochondria) was kept in the stock by crossing females with *Wolbachia* with indicated males. The mitochondria are only shown in females because of its strictly maternal transmission. All males were free of *Wolbachia* infection. Dashed lines indicate the genotype selected from the previous cross. Virgin female in the first cross were considered mutant in all chromosomes (*), for illustrative purposes. Question marks (?) represent recombined chromosomes.

**S5 Fig.**
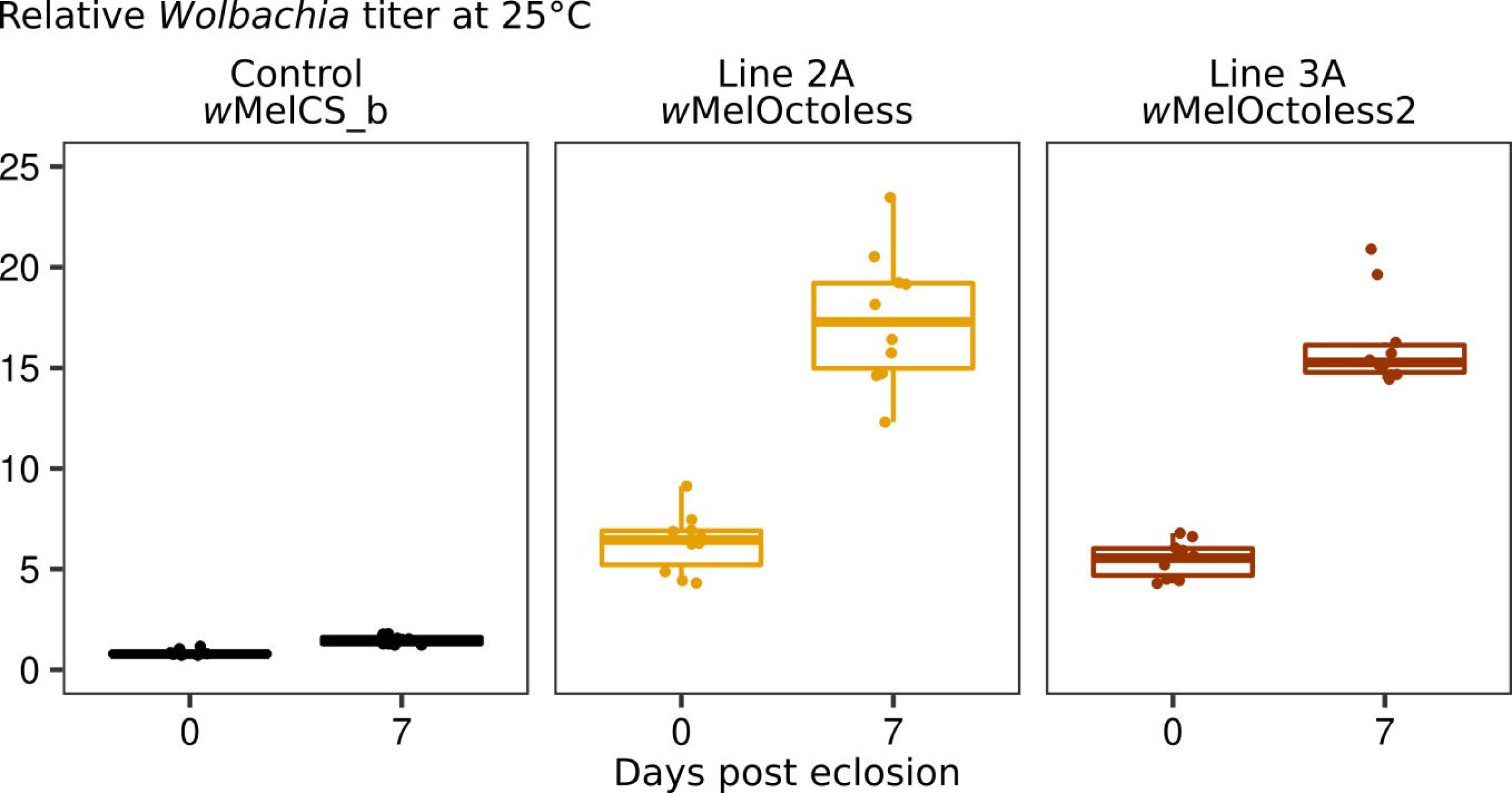
Proliferation of *w*MelOctoless and *w*MelOctoless2 in a host isogenic genetic background. Relative *Wolbachia* titres in *D. malanogaster* males carrying *w*MelOctoless and *w*MelOctoless2 at 0 and 7 days post adult eclosion, at 25°C. This experiment was set-up as described in Fig 1. Relative *Wolbachia* titre was determined using qPCR and normalized to that of 0-1 days-old *w*MelCS_b-infected males. Each dot represents the relative titre of a single male.

**S6 Fig.**
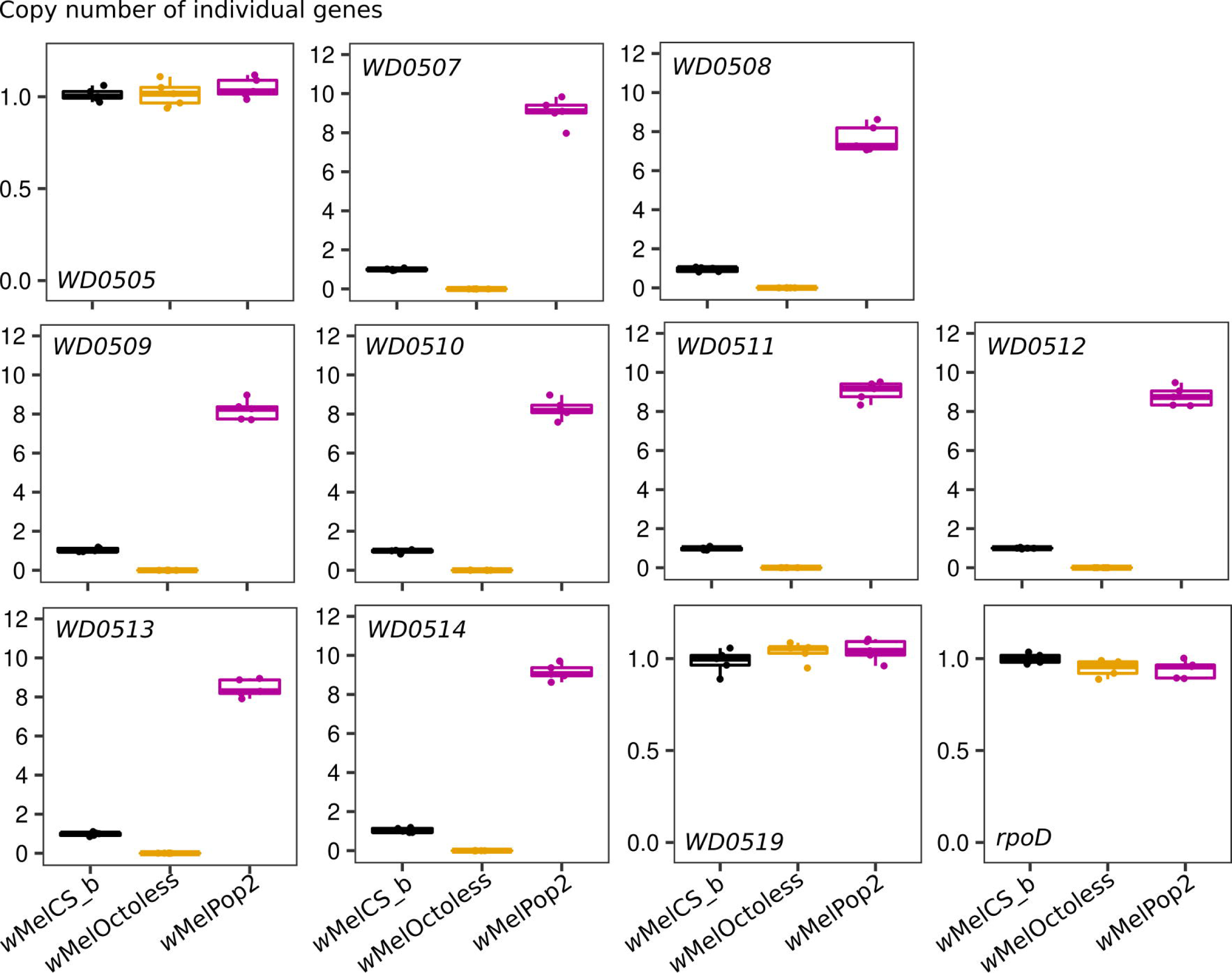
Confirmation of amplification and deletion of Octomom genes by qPCR. The amplification and deletion of individual Octomom genes (wMel loci *WD0507–WD0514*) was confirmed using qPCR in *w*MelPop2 and *w*MelOctoless, respectively. The copy number of three genes outside the Octomom region (wMel loci *WD0505, WD0519*, and *rpoD*) were also determined. Five females carrying *w*MelCS_b, *w*MelPop2, and *w*MelOctoless were used in the analysis. The copy number of *w*MelPop2 and *w*MelOctoless genes is relative to that of *w*MelCS_b.

**S7 Fig.**
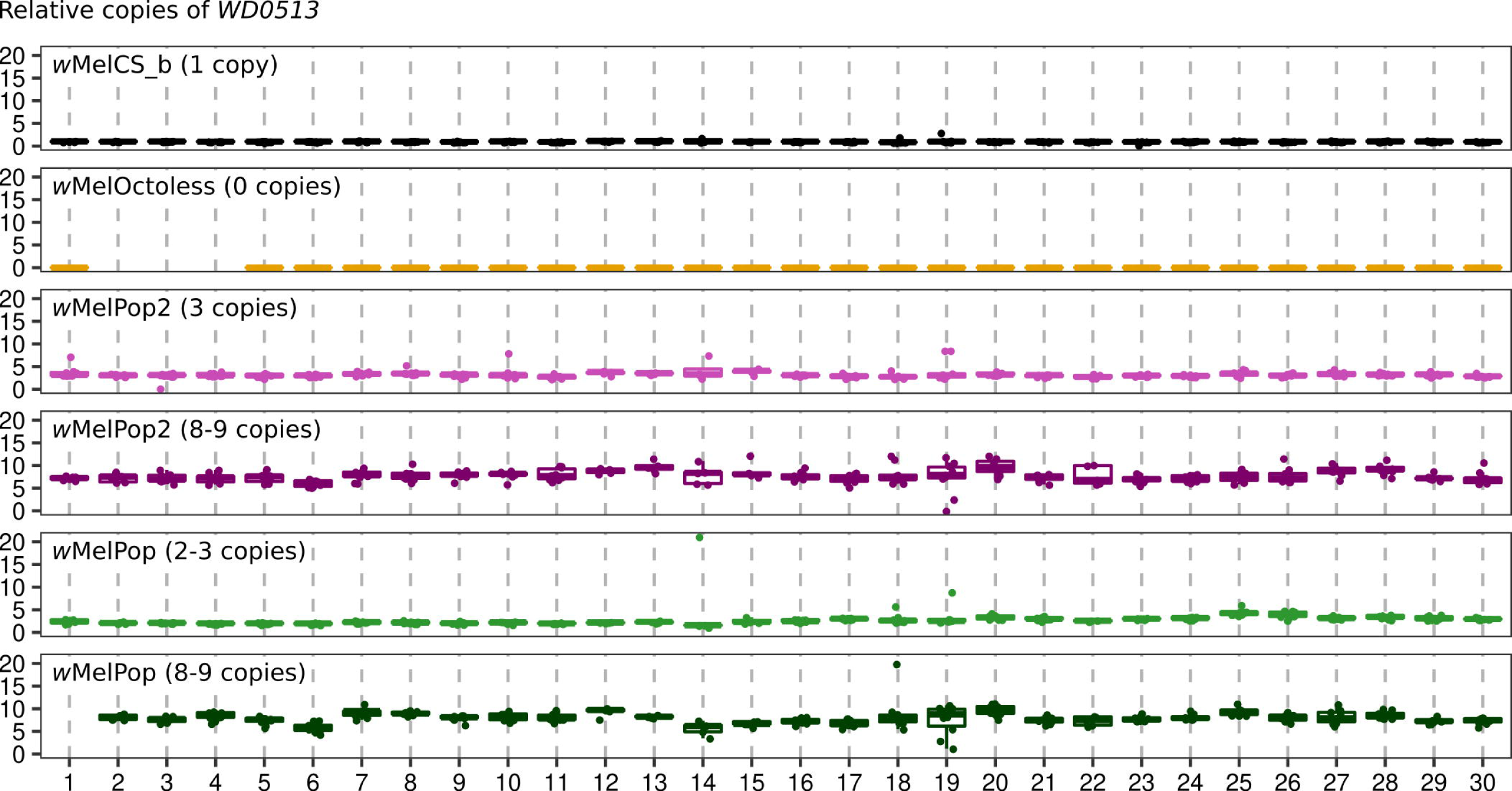
Selection for lines carrying *Wolbachia* with a desired Octomom copy number. The relative copy number of genomic *WD0513* in *Wolbachia*-carrying stocks throughout 30 fly generations. Each generation, 5-20 females were randomly collected for egg-laying for 3-4 days and used to determine the relative copy number of *WD0513*, as a proxy for the Octomom copy number. The progeny of a single female was used to set up the next generation. qPCR results were normalized to that of *w*MelCS_b, which has a single copy of Octomom per genome.

**S8 Fig.**
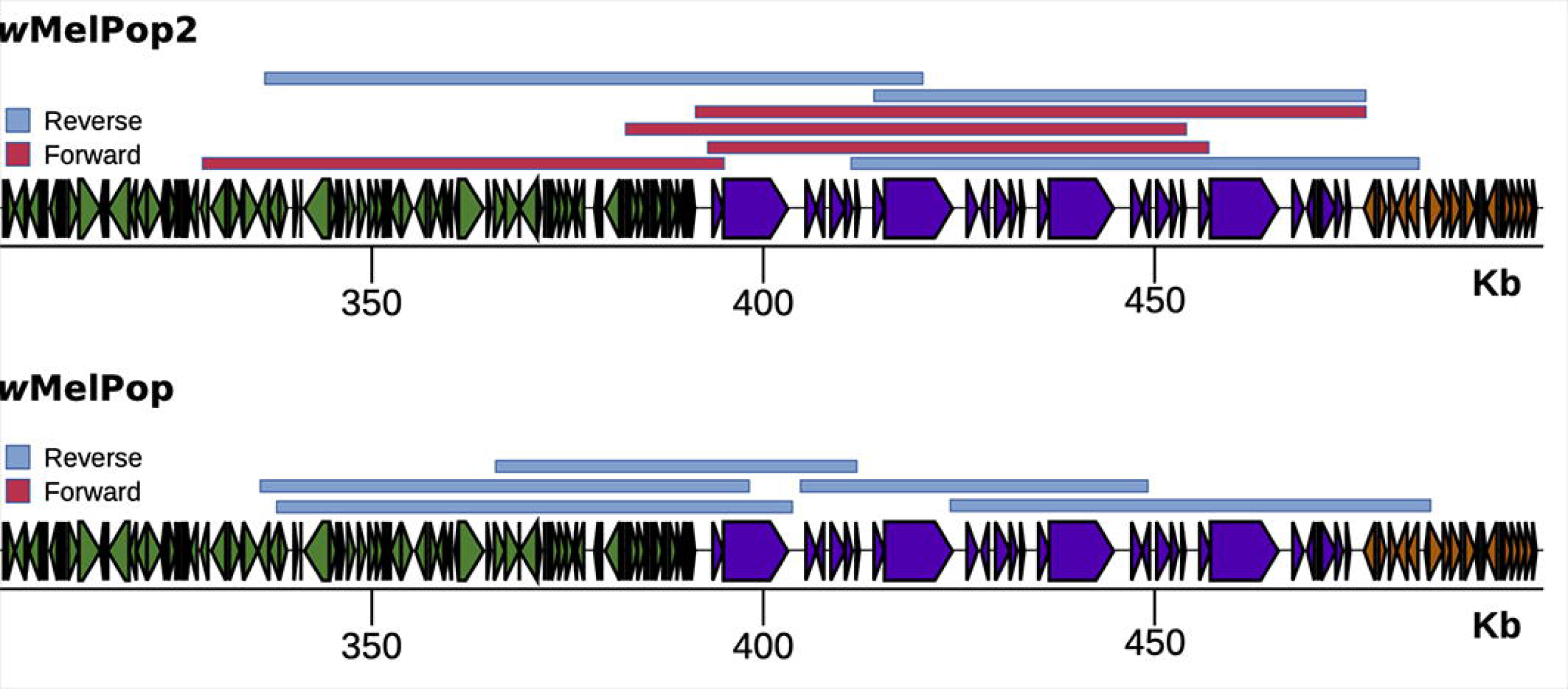
Octomom region is amplified in tandem in *w*MelPop2 and *w*MelPop. Oxford Nanopore MinION reads supporting the amplification of the Octomom region in tandem in *w*MelPop2 and *w*MelPop *Wolbachia* variants. We mapped *w*MelPop2 and *w*MelPop long reads (BioProject: PRJNA587443) to the the Octomom region in their genomes (Accessions CP046922.1 and CP046921.1, respectively) using minimap2 v2.17-r941 [48] and plotted the alignment summary (S7 Table) for illustrative purposes.

**S9 Fig.**
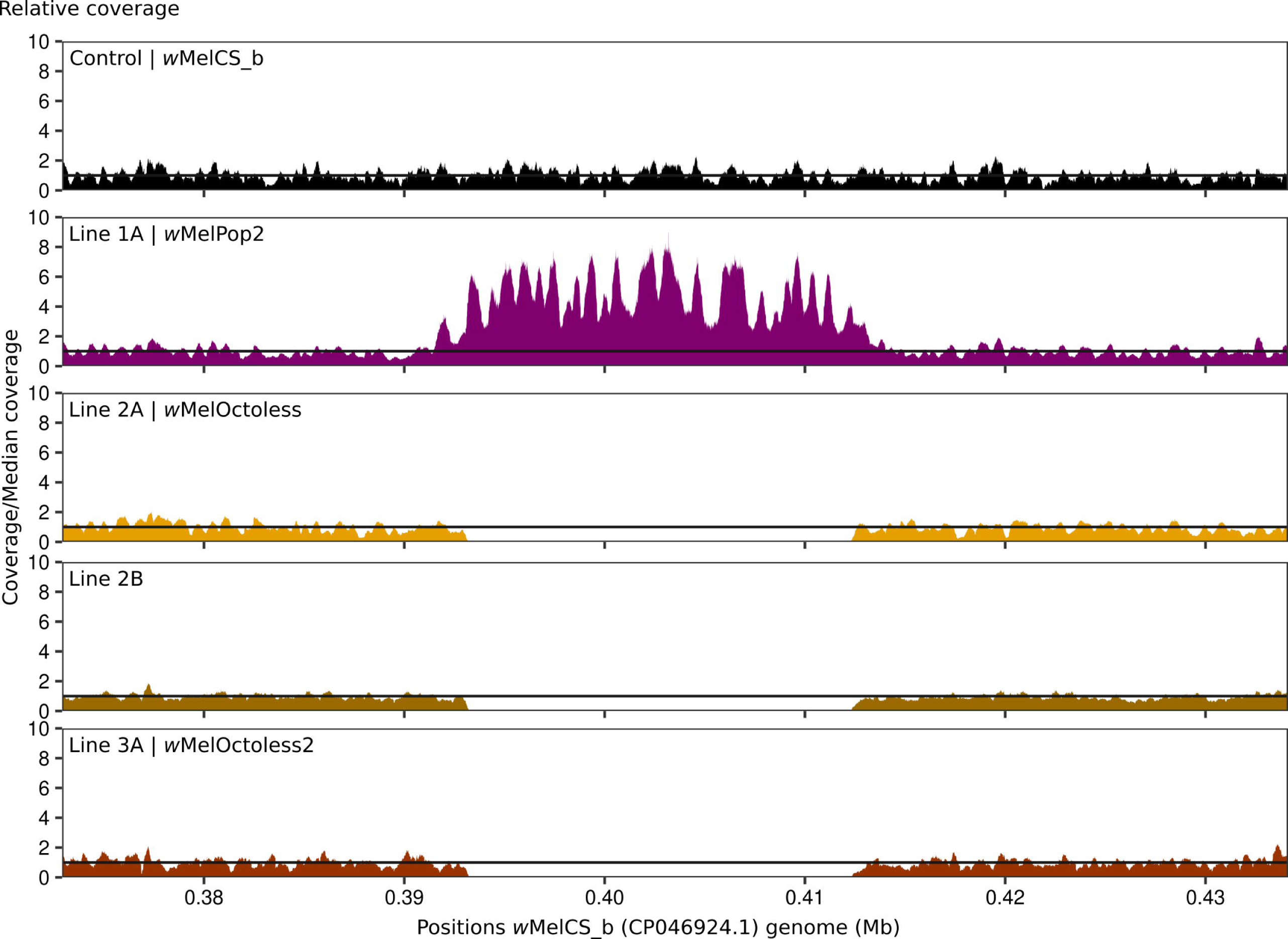
Identification of the genetic bases for over-proliferation of the *Wolbachia* in Line 2B and Line 3A (wMelOctoless2). Relative coverage in the genomic region containing the Octomom region. As in Fig 2B, Illumina paired-end reads were mapped to *w*MelCS_b (GenBank: CP046924.1) genome, and the number of reads mapping to each position were normalized by dividing to the median coverage across the genome. Coverage information for *w*MelCS_b, *w*MelPop2 and *w*MelOctoless is also given in Fig 2B. We identified the deletion of Octomom as the cause of proliferation in lines 2B and line 3A (wMelOctoless2), as no other difference was found when compared to *w*MelCS_b.

**S10 Fig.**
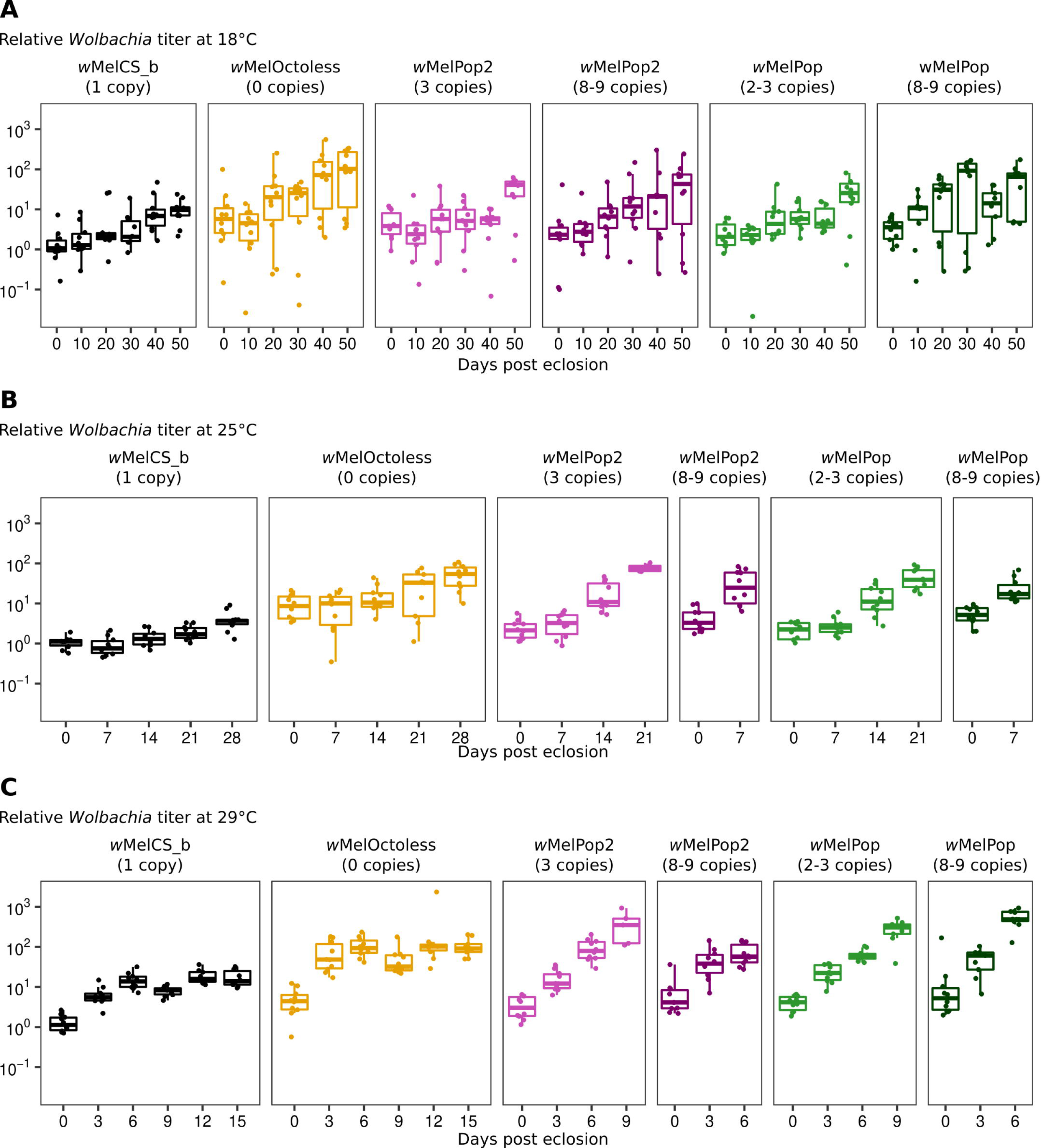
The amplification or deletion of Octomom increase *Wolbachia* proliferation rate in adults. Time-course of relative *Wolbachia* titres in adults at 18°C (A), 25°C (B) and 29°C (C) with different *Wolbachia* variants. Replicate of experiment shown in Fig 3. *Wolbachia* titres were determined and analysed as described for Fig 3.

**S11 Fig.**
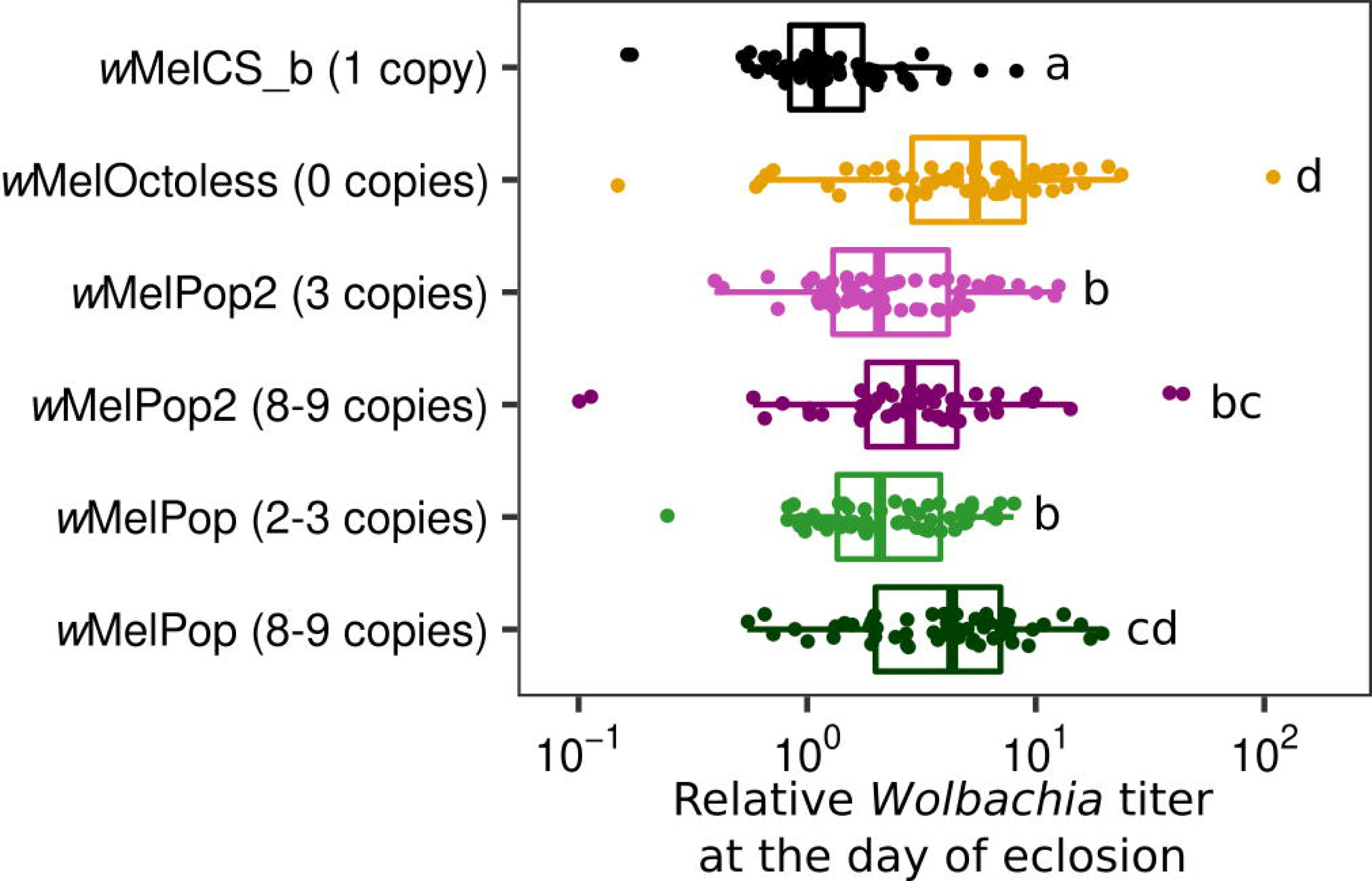
Octomom copy number determines *Wolbachia* titres on the day of adult eclosion. Relative *Wolbachia* titres on the day of adults eclosion. Males developed at 25°C were collected within 24 hours after eclosion for *Wolbachia* titre measurement using qPCR. Data used in this figure are also shown in Fig 3 and S10 Fig (time point 0). Letters represent significant groups after p-value correction.

**S12 Fig.**
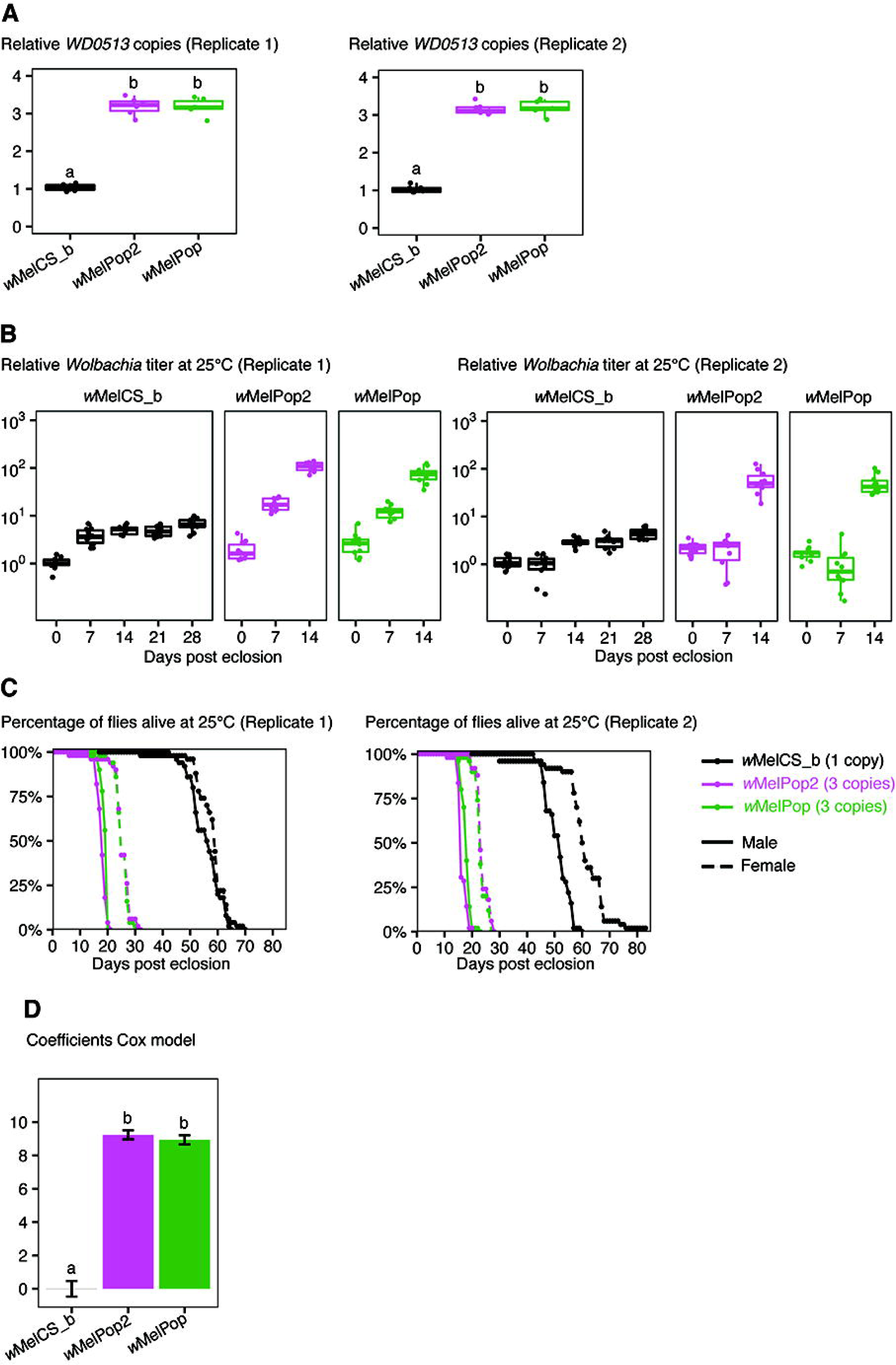
wMelPop2 and *w*MelPop are phenotypically indistinguishable. (A) *WD0513* copy number of *w*MelPop2 and *w*MelPop in two experimental replicates. Using *WD0513* as a proxy, the Octomom copy number of *w*MelPop2 and *w*MelPop was tightly controlled prior to phenotypic comparison. (B) *Wolbachia* relative titres at 25°C. The progeny of *w*MelPop2- and *w*MelPop-infected females carrying three copies of Octomom was used to set up the experiments. Males that developed at 25°C were collected upon eclosion, aged to specific time-points and used to determine *Wolbachia* titres using qPCR. *Wolbachia* titres were normalized to that of *w*MelCS_b-carrying flies collected on the day of eclosion. Proliferation rates of *w*MelPop2 and *w*MelPop were not different (p = 0.32). (C) Lifespan of males (solid lines) and females (dashed lines) flies at 25°C. Males were transferred to new vials every five days, while females every four days. (D) Coefficients of a Cox mixed model, representing the effect of *w*MelPop2 and *w*MelPop on the lifespan relative to *w*MelCS_b-carrying flies. *w*MelPop2 and *w*MelPop was equally pathogenic (p = 0.29).

**S13 Fig.**
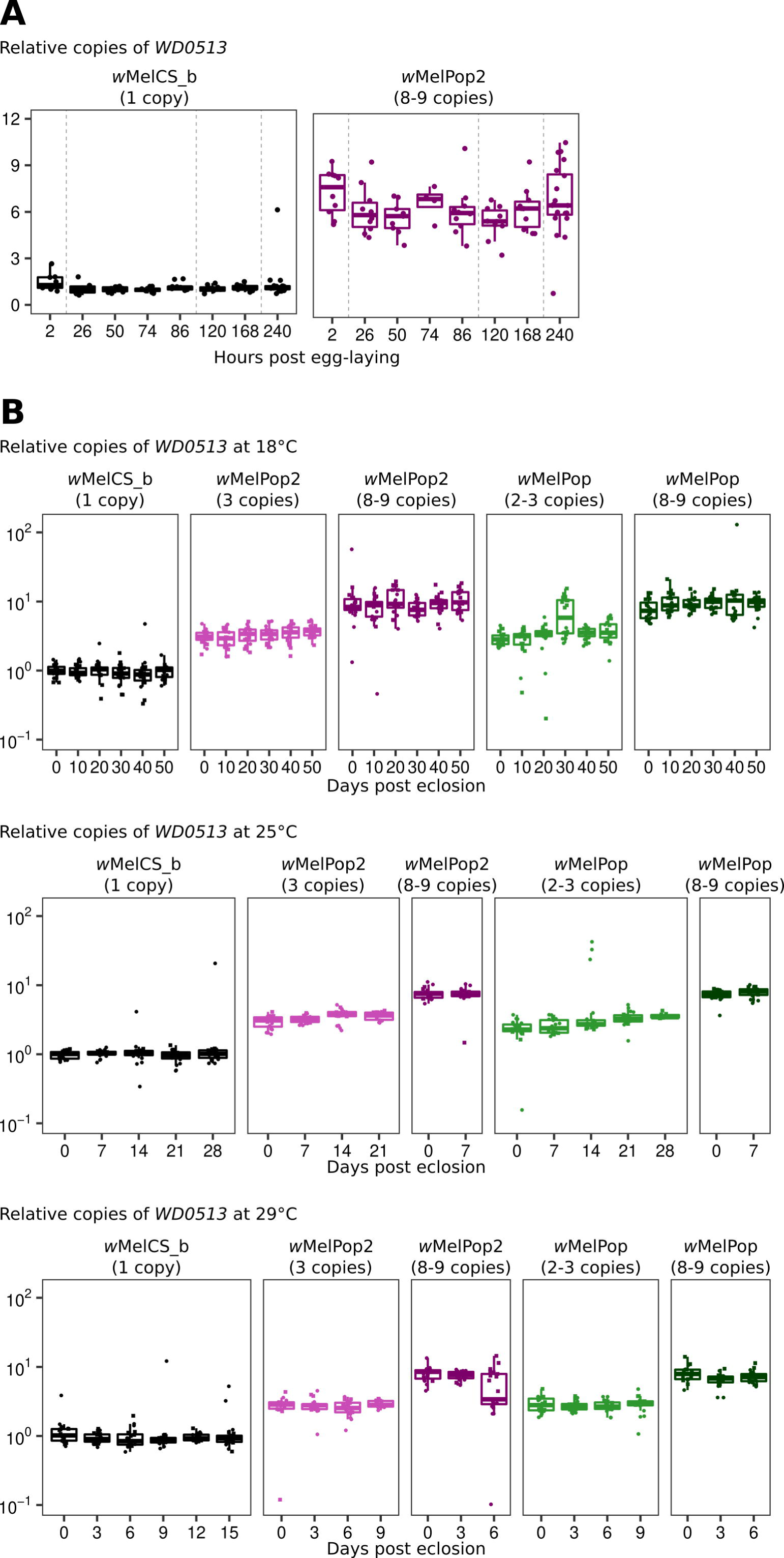
Octomom copy number dynamics throughout fly development and during adult life. Relative copies of *WD0513* throughout *D. melanogaster* development (A) and during adult life (B). *WD0513* relative copy numbers were determined in samples shown in Fig 4 (for panel A) and Fig 3 and S10 Fig (for panel B). *WD0513* copies were normalized to that of 0-1 old *w*MelCS_b-infected males. (A) Vertical dashed lines separate developmental stages (i.e. eggs, larvae, pupae, and adults). The x-axis is not in a continuous scale. (B) The two replicates are represented by different symbols.

**S14 Fig.**
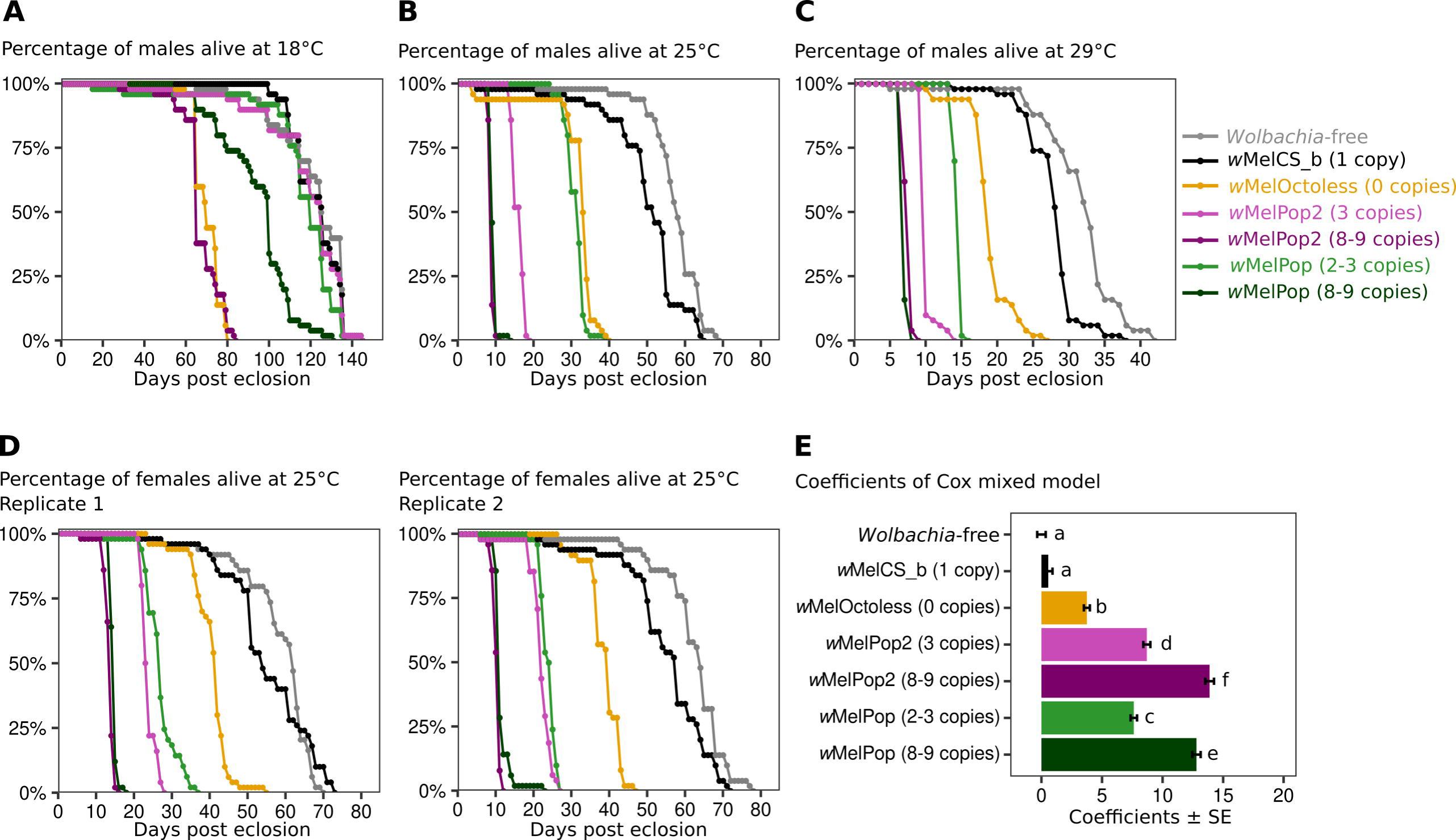
wMelPop2 and *w*MelOctoless are pathogenic to both males and females. Lifespan of *D. melanogaster* males at 18°C (A), 25°C (B), and 29°C (C). Survivorship was determined as in Fig 5. This is a replicate of Fig 5 (D) Survival of *D. melanogaster* females at 25°C. Survival was determined as in Fig 5, except that females were transferred to new vials every four days. The experiment was performed twice. (E) Coefficients of a Cox mixed model of the lifespan of females relative to *Wolbachia*-free control. Both replicate experiments were pooled for statistical comparisons. Bars represent the standard error of the coefficient, and letters the statistically significant groups.

**S15 Fig.**
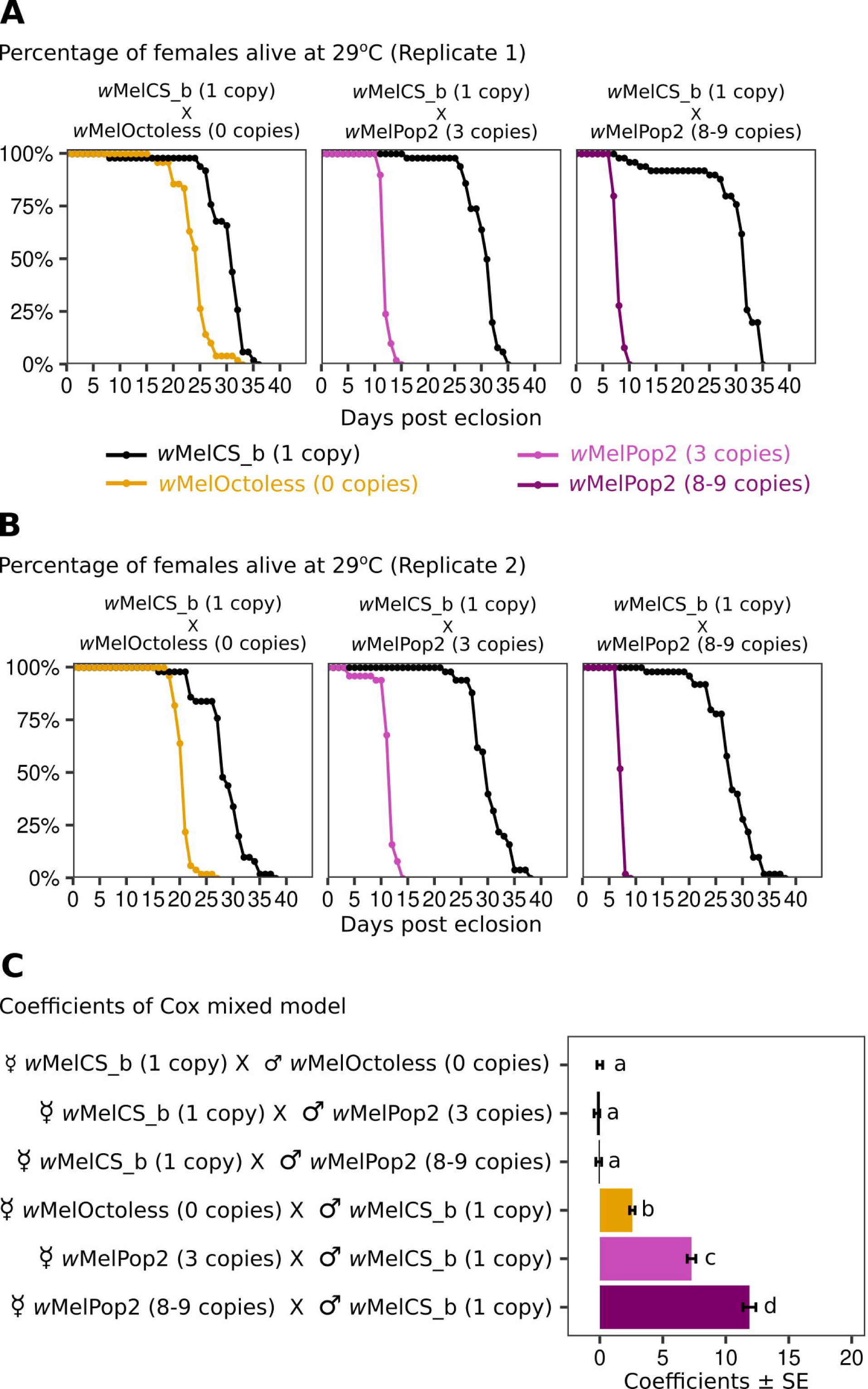
Wolbachia variants, not differences in the host genetic background, are pathogenic. (A-B) Survival of *D. melanogaster* females at 29°C. Virgin *w*MelCS_b-carrying females were crossed with males carrying *w*MelOctoless or *w*MelPop2 (with 3 or 8-9 Octomom copies) and vice-versa. The resulting progeny developed at 25°C and was placed at 29°C after adult eclosion. The survival of 50 female progeny, which have the same genetic background but differ in *Wolbachia* infection, was determined per condition, per replicate. Females were maintained in groups of ten and transferred to new vial every four days. The experiment was performed twice. (C) Coefficients of a Cox mixed model representing the effect of the parental crosses on the survivorship of females. Significance was accessed after p-value correction for multiple comparisons, and significant groups are represented by letters.

**S16 Fig.**
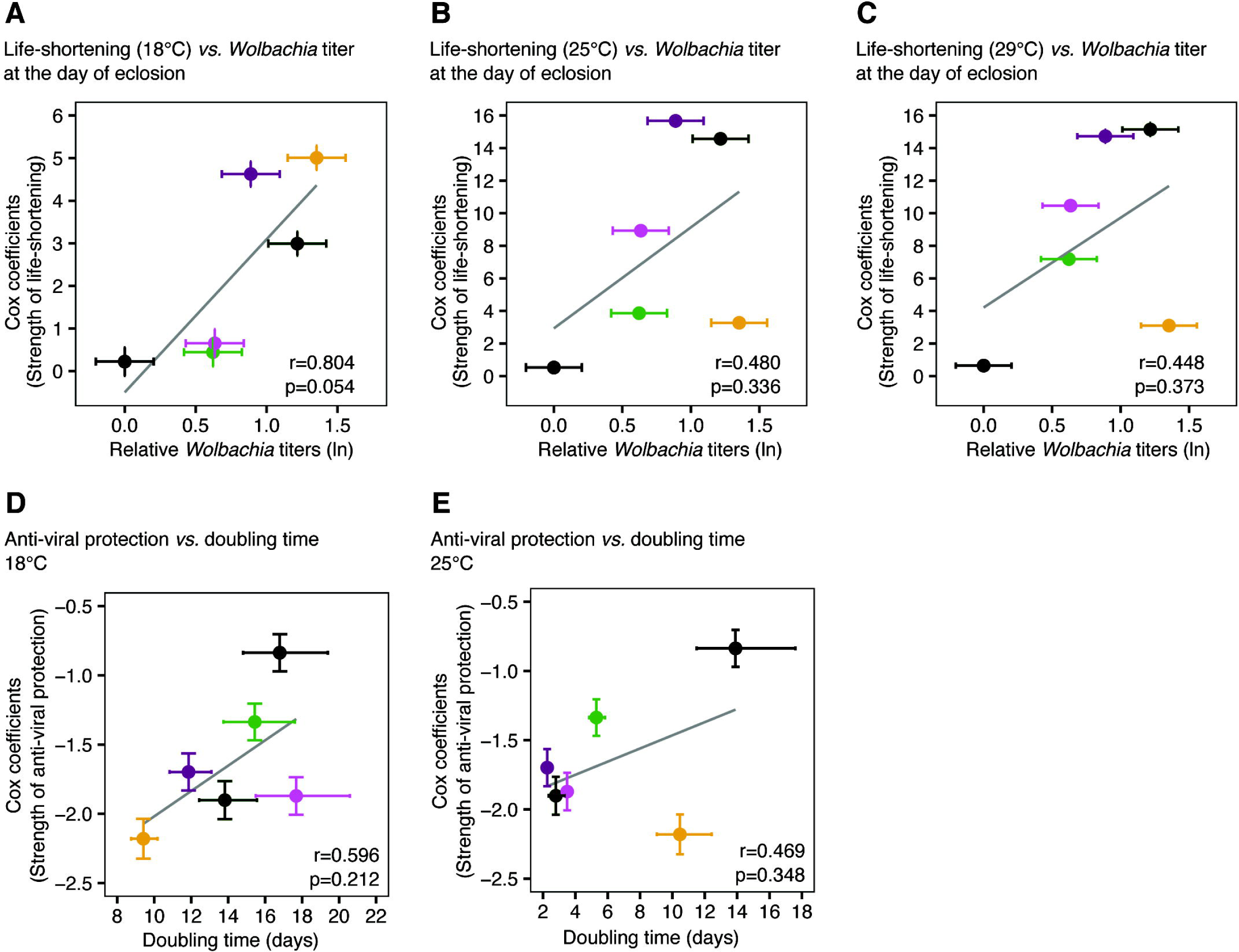
Correlation between *Wolbachia*-induced phenotypes and bacterial titres or doubling time. (A-C) Correlation between *Wolbachia* titre at the day of eclosion and the strength of life-shortening phenotype determined at 18°C (A), 25°C (B), and 29°C (C). The y-axis represents the strength of *Wolbachia* life-shortening phenotype (estimated using Cox mixed model shown in Fig 5). The x-axis represents the natural log of the relative *Wolbachia* titre estimated using a linear mixed model. Bacterial titres were normalized to that of *w*MelCS_b-infected flies (shown in S11 Fig). (D and E) The correlation between the strength of anti-viral protection and *Wolbachia* doubling time. The y-axis represents the strength of anti-viral protection (estimated using Cox mixed model shown in Fig 6). The x-axis represents *Wolbachia* doubling time in adults at 18°C (D), or 25°C (E) (shown in Table 1). The Pearson correlation coefficient (r) and its significance (p) are given in each panel. A grey line represents the trend (fit of linear regression). Error bars represent the standard errors of the estimates. None of these correlations were statistically significant and they complement correlations shown in Fig 5 and Fig 6.

**S17 Fig.**
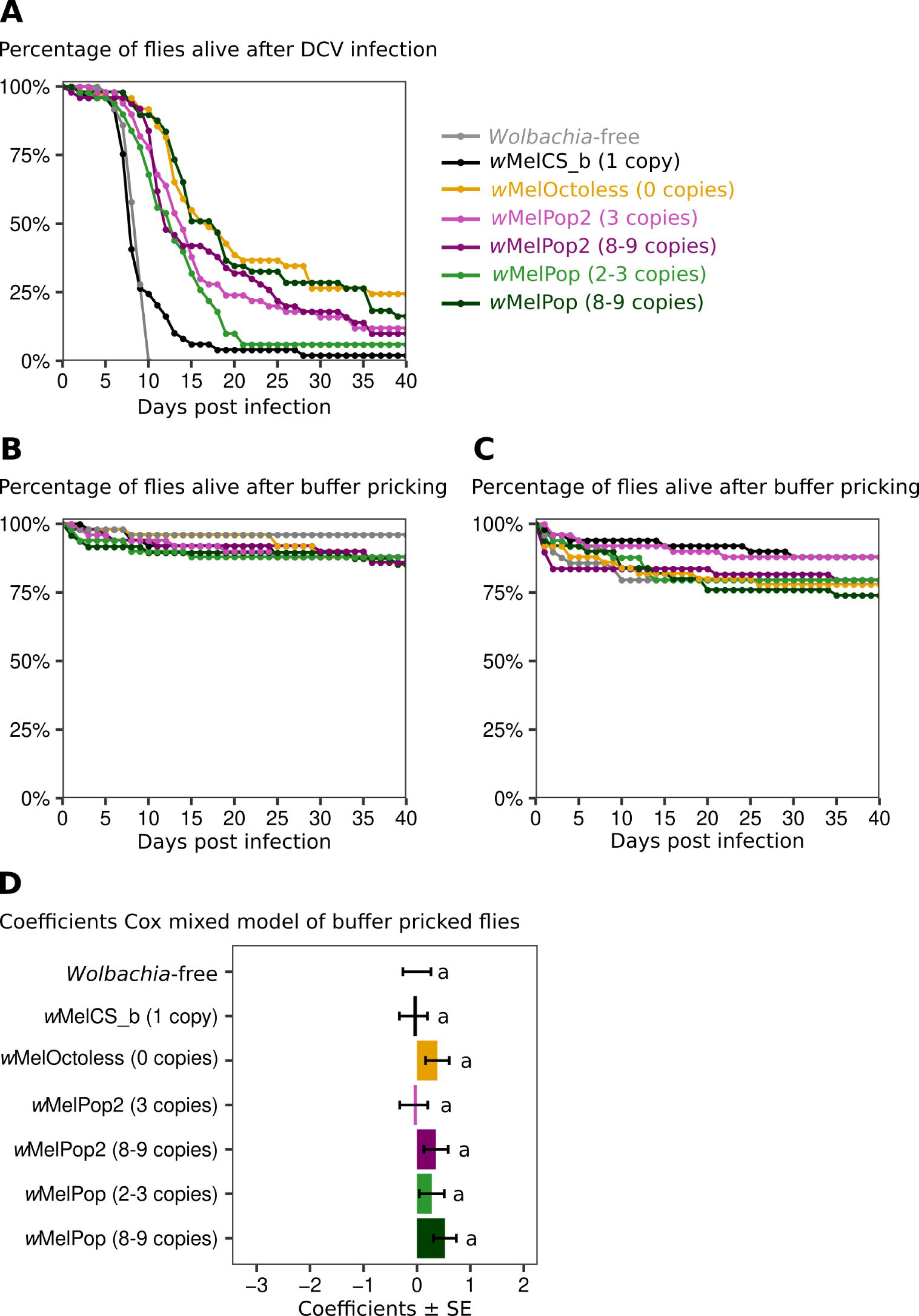
Survival of flies with different *Wolbachia* variants after challenge with DCV or buffer solution. (A) Survival of males carrying different *Wolbachia* variants after a challenge with DCV (A) or a buffer solution (B and C). Fifty 3-5 days-old Drosophila males, per line, were pricked with DCV (10^9^ TCID_50_/ml) or buffer and survival curves were determined at 18°C for 40 days. A is a replicate of Fig 6A, B and C are controls for these experiments. (D) Coefficients of Cox mixed models of buffer-pricked flies. Both replicates were pooled for statistical analysis. Bars represent the standard error of the estimate, and the letters the statistically significant groups after p-value correction.

**S1 Table. Number of F1 females screened for new over-proliferative *Wolbachia* variants per experimental condition.**

*w*MelCS_b-infected G0 females, raised in control or antibiotic-treated food (12.5 μg/ml), were fed different doses of ethyl-methanesulfonate (EMS) and allowed to lay eggs in individual vials. F1 females were collected as virgins, mated to non-mutagenized males and also allowed to lay eggs individually. F1 females were used for *Wolbachia* titre measurement when were 10-days old. Number of F1 females tested per experimental condition is shown.

S2 **Table. Coverage statistics of the sequencing project.**

Coverage statistics (mean and range) of Illumina reads mapped to either *Wolbachia* or mitochondria of *D. melanogaster* Release 6 genome sequence (KJ947872.2:1–14,000). Sequencing data of each *Wolbachia* variants are mapped to own genome assembly (BioProject ID: PRJNA587443), except for *Wolbachia* in Line 2B and *w*MelOctoless2 which were mapped to *w*MelCS_b genome (Accession: CP046924.1). ND – not determined.

**S3 Table. Flies infected with new over-proliferative *Wolbachia* variants did not inherit mutated mitochondria.**

Illumina reads on flies infected with different *Wolbachia* variants were mapped to the mitochondria of *D. melanogaster* Release 6 genome sequence (KJ947872.2:1–14,000). A summary of the mapping is given in S2 Table. The mitogenome of flies infected with *w*MelCS_b, *w*MelOctoless and *w*MelPop2 was identical. We found an SNP unique to flies infected with *w*MelCS-like variants (G→A on position 10,793) but absent in flies infected with *w*MelPop. We confirmed this SNP using Sanger sequencing.

**S4 Table. Assembly and annotation statistics.**

*Wolbachia* genomes were assembled using the Unicycler v0.4.8-beta pipeline and annotated using NCBI Prokaryotic Genome Annotation Pipeline v4.10. *w*Mel reference genome (Accession: AE017196.1) is included for comparison purposes.

**S5 Table. SNPs and indels between newly assembled *w*Mel and *w*Mel reference genome.**

The genome of a newly assembled Cluster III *w*Mel *Wolbachia* variant (Accession: CP046925.1) was aligned to *w*Mel reference genome (Accession: AE017196.1) using Mauve v2.4.0. All the differences were confirmed via Sanger sequencing.

**S6 Table. SNPs and indels between *w*MelCS_b and and *w*Mel reference genome.**

The genome of *w*MelCS_b (Accession: CP046924.1) was aligned to *w*Mel reference genome (Accession: AE017196.1) using Mauve v2.4.0.

**S7 Table. Alignment summary of long reads supporting the amplification of the Octomom region in tandem.**

Long reads (MinION, Oxford Nanopore) reads supporting the amplification of the Octomom region in tandem in *w*MelPop2 (Accession: CP046922.1) and *w*MelPop (Accession: CP046921.1) genomes. Long reads were mapped to Octomom region using minimap2 v2.17-r941 and the number of Octomom copies determined using blastn v2.8.1+.

**S8 Table. Primers used for amplification and quantification of individual *Wolbachia* genes.**

Primers used in this study have been previously described (Chrostek 2013 and Chrostek 2015).

**S9 Table. List of primers used to improve *Wolbachia* draft genomes.**

Primers used to amplify and sequence, using Sanger technology, genomic regions containing predicted differences between *Wolbachia* draft genomes.

**S1 Text. Confirmation of the amplification and deletion of the Octomom in *w*MelPop2 and *w*MelOctoless, respectively.**

The genomes of *w*MelCS_b, *w*MelPop2 and *w*MelOctoless were aligned using Mauve v2.4.0. The three-fold amplification of Octomom in *w*MelPop2 and its deletion in *w*MelOctoless were the only difference identified when compared with *w*MelCS_b.

